# Camera traps unable to determine whether plasticine models of caterpillars reliably measure bird predation

**DOI:** 10.1101/2024.10.01.616075

**Authors:** Laura Schillé, Nattan Plat, Luc Barbaro, Hervé Jactel, Frédéric Raspail, Jean-Baptiste Rivoal, Bastien Castagneyrol, Anna Mrazova

**Affiliations:** BIOGECO, INRAE, University Bordeaux, Cestas, France; DYNAFOR, University of Toulouse, INRAE, Toulouse, France; Institute of Entomology, Biology Centre of the Czech Academy of Sciences, Ceske Budejovice, Czech Republic

**Author notes:** These authors contributed equally to this work (Laura Schillé, Nattan Plat). These authors also contributed equally to this work (Bastien Castagneyrol, Anna Mrazova). **Contributions** A.M., L.S., and N.P. developed the original idea. A.M., B.C., L.S., and N.P. developed the methodology. A.M., B.C., H.J., JB.R., L.B., L.S., and N.P. conducted data collection. F.R. implemented the computer code. A.M. and L.S. did the formal analysis and wrote the original draft, and all co-authors provided critical reviews of the manuscript.

**Keywords:** artificial prey, bird predation, camera trap, ecological methods, fake prey, plasticine caterpillar models, real caterpillars

## Abstract

Sampling methods that are both scientifically rigorous and ethical are cornerstones of any experimental biological research. Since its introduction 30 years ago, the method of using plasticine prey to quantify predation pressure has become increasingly popular in biology. However, recent studies have questioned the accuracy of the method, suggesting that misinterpretation of predator bite marks and the artificiality of the models may bias the results. Yet, bias *per se* might not be a methodological issue as soon as its statistical distribution in the samples is even, quantifiable, and thus correctable in quantitative analyses. In this study, we focus on avian predation of lepidopteran larvae models, which is one of the most extensively studied predator-prey interactions across diverse ecosystems worldwide. We compared bird predation on plasticine caterpillar models to that on dead caterpillars of similar size and color, using camera traps to assess actual predation events and to evaluate observer accuracy in identifying predation marks *a posteriori*. The question of whether plasticine models reliably measure insectivorous bird predation remained unanswered, for two reasons: (1) even the evaluation of experienced observers in the posterior assessment of predation marks on plasticine models was subjective to some extent, and (2) camera traps failed to reflect predation rates as assessed by observers, partly because they could only record evidence of bird presence rather than actual predation events. Camera traps detected more evidence of bird presence than predation clues on plasticine models, suggesting that fake prey may underestimate the foraging activity of avian insectivores. The evaluation of avian predation on real caterpillar corpses was probably also compromised by losses to other predators, likely ants. Given the uncertainties and limitations revealed by this study, and in the current absence of more effective monitoring methods, it remains simpler, more cost-effective, ethical, and reliable to keep using plasticine models to assess avian predation. However, it is important to continue developing improved monitoring technologies to better evaluate and refine these methods in order to advance research in this field.

## Introduction

The functional role of insectivorous birds in terrestrial ecosystems (1,2) has been the focus of many studies aimed at understanding predation patterns, feeding behavior, predator control of prey populations, and evolutionary ecology (3–6). Traditional methodological approaches to these research topics include direct field observations (7), experimental manipulation in the laboratory (8,9) or in the field (10), and gut or fecal DNA analyses (11,12). Each of these methods has its own set of limitations regarding standardization, logistics, and ethical considerations such as not sparing live prey or handling captive birds. To address these challenges, the use of artificial prey, particularly those made of plasticine, introduced over 30 years ago by Brodie III (13), has become a valuable tool in bird predation research (14). The method is primarily appreciated for its ability to preserve the bite marks of predators, the identification and quantification of which has been used to infer attacks and predation risk from specific predators.

While plasticine caterpillar models are widely employed as simple, standardized prey models to investigate predation activity in diverse ecosystems (15–17), recent studies have raised concerns regarding their limitations and potential inaccuracies (14,18–20). The main concerns relate to the accuracy of inferences of predator identity and activity from the posterior assessment of putative predation marks on prey models.

### Inaccuracy of predator identity

Observers may struggle to attribute a considerable portion of attack marks to a specific predator type, e.g., bird, reptile, or arthropod (18,19). For example, Valdés-Correcher et al. (21) found that untrained scientists misidentified up to 26% of predation mark on photos of attacked plasticine caterpillar models; while experienced scientists misidentified 21% of the marks. These findings were based on comparisons to evaluations conducted by an expert who examined the actual plasticine caterpillar models. They underscore the difficulty of accurately identifying predator marks, particularly when only photographs are available. This limitation can introduce observer bias, especially among untrained observers who may face even greater challenges. The lack of a standardized training system further compounds this variability, as it prevents observers from gaining knowledge in recognizing such marks consistently (18,22; but see Low et al. (23), for examples of predator-type classified bite mark images). Consequently, there may be differences of interpretation between researchers.

### Inaccuracy in determining real predation events among all identified marks

Even though marks were attributed to birds, questions would remain about what one can infer from them. Incidental contact or non-predatory interactions (e.g., by bird claws, attraction of non-predatory animals, or curiosity) with the plasticine caterpillar models can result in marks that are difficult to distinguish from genuine predation attempts, potentially leading to an overestimation of predation rates (24). The misinterpretation of marks can lead to distorted results on predation pressure and misleading conclusions about predator-prey interactions, especially in areas where the attack rate is low (25).

Furthermore, the artificial appearance and texture of plasticine may not adequately simulate the characteristics of real prey, including smell, possibly affecting the behavior of insectivorous birds and thus the validity of predation assessment (19). The literature contains several studies that compared the attack rates on plasticine larvae to those on real larvae, with the latter being quantified based on missing or partially consumed prey items (19,20,24). As opposed to plasticine caterpillar models, the attack rate on live and dead prey were similar (19), thus supporting the use of dead prey as a more ethical alternative to live prey (to prevent gluing or pinning them while alive) when comparing real and plasticine prey. By simultaneously studying predation on dead caterpillars versus plasticine models, it becomes possible to compare the response of predators to both types of prey within the same experimental design with the use of passive monitoring methods. This comparative approach makes it possible to test the assumptions on which the plasticine caterpillar model method is biased and to correct it in relation to observable predation patterns on real dead prey. Quantifying this bias is critical to determining whether plasticine models can reliably mimic real prey, and thus whether the data they provide can reflect an accurate depiction of predation pressure.

Studies comparing predation on both real caterpillars and plasticine models have shown discrepancies, revealing that plasticine models may not be a sufficiently accurate method for assessing predation (19), particularly across large biogeographical ranges (24), varying thermal regimes (26), different seasons of the bird life cycle (20), and for different taxa (19). There appears to be a gap in the literature regarding more detailed analyses that elucidate the origins of the marks found on plasticine models. Moreover, missing plasticine models are usually excluded from the statistical analyses (16,27). Yet, at least part of those missing models can be the result of a predation attempt that was not recorded. On the contrary, studies working with real prey have often assumed that a missing prey was equivalent to a depredated prey (19,20,24), an assumption that may not always hold true. Without direct observation, the reasons behind the disappearance of the prey remain speculative.

To address these methodological uncertainties, and following recommendations from Bateman et al. (18) and Rößler et al. (22), we used camera traps. Since camera traps had previously been employed to study bird frugivory (28,29), we suggested that their use could also enable the recording predation on plasticine models (hereafter “models”) paired with dead caterpillars (hereafter “corpses”) on the experimental trees. We hypothesized that using camera traps could visually confirm predation attempts, providing a direct and unbiased record of predator interactions with the prey. This visual evidence could enable the differentiation between actual predation marks and those resulting from incidental contact or environmental factors (e.g., rubbing by leaves or branches). Ultimately, our goal was to use camera traps to efficiently compare bird predation on corpses vs. on models and to evaluate posterior observer assessment of predation marks. We believed that the results of this study would help refine the plasticine caterpillar model method as a tool for ecological research, ensuring that it produces scientifically relevant results.

We used camera traps as a tool to test the reliability of the plasticine caterpillar method by quantifying predation events on both prey types and possible sources of bias in inferences made from plasticine models. We first assessed observer-related bias in the detection of predation clues on plasticine models, hypothesizing that experienced observers would show a high degree of consistency when identifying bird predation marks on these models (H1). We further hypothesized that observer detection of predation clues would lead to an overestimation of predation on plasticine models by attributing unrelated or uncertain marks to actual predation attempts (H2). Taking into account the innate ability of insectivorous birds to recognize suitable prey and the differences in the surface texture of real vs. plasticine caterpillars, we hypothesized that there would be evidence of more predation on corpses than on models, both assessed using camera traps (H3). Finally, we hypothesized that observer detection of predation would lead to an overestimation of the predation on corpses by accounting for missing individuals as a bird predation event, lacking direct evidence for why the prey disappeared (H4). If we obtained comparable results of the bias between observers and cameras for both models and corpses and if the test of H3 provided evidence of more predation on corpses than on models, we could conclude that the use of corpses with human post-detection might be a more accurate method than the use of plasticine models with human post-detection. The present article was initially evaluated as a registered report (30). For detailed information on all scientific questions, hypotheses, predictions, and analysis plans, see S1 Table, S2 and S3 Figs.

## Material and Methods

### Study area

We carried out this experiment in 2024 in a pine and oak forest stand of 5ha near Bordeaux, France (44°440N, 00°460W). The climate is oceanic, with 12.8°C average temperature and 870mm annual rainfall. We synchronized the start of the experiment as closely as possible with the breeding period of local insectivorous birds, specifically, when the chicks were in the nest and fed by the parents (starting generally mid-April for resident breeders or early migrants). We choose this timing to quantify bird predation by experienced adults rather than later exploratory predation attempts by young, inexperienced birds. The main insectivorous bird species in the study area were Great Tit (*Parus major*), Eurasian Blue Tit (*Cyanistes caeruleus*), Eurasian Blackcap (*Sylvia Atricapilla*), Common Redstart (*Phoenicurus phoenicurus*), Eurasian Blackbird (*Turdus merula*), Western Bonelli’s Warbler (*Phylloscopus bonelli*), Common Chiffchaff (*Phylloscopus collybita*), Short-toed Treecreeper (*Certhia brachydactyla*), Eurasian Nuthatch (*Sitta europaea*), European Robin (*Erithacus rubecula*), Eurasian Wren (*Troglodytes troglodytes*), and Great Spotted Woodpecker (*Dendrocopos major*) (31).

These bird species have home ranges that vary between 0.3 ha for the Blue Tit (32) and 7.9 ha for the Common Chiffchaff (33), the latter value being extreme as compared to most of the insectivorous species (34,35). It is therefore likely that multiple individuals of the same insectivorous bird species defended separate territories in the study area. Additionally, we distributed the experiment across the whole forest stand (see below), where oak trees were consistently observed to host a substantial number of herbivorous insects (N. Plat, A. Mrazova, and L. Schillé personal observation). A sufficient abundance of natural prey reduces the risk of birds altering their typical feeding behaviors and minimizes overly frequent encounters with artificial prey items. Therefore, we believe it is unlikely that the birds learned to avoid the corpses and models used during the experiment.

### Study design

First, we haphazardly selected 12 pedunculate oaks (*Quercus robur*, the dominant broadleaved species of the area) located at least 10m apart from each other. Due to practical constraints, we chose branches that were accessible from the ground to attach prey items, which prevented us keeping the same branch orientation between trees. This is also the approach that has been used in a previous study evaluating the plasticine prey method (20). On one branch of each tree, we glued five plasticine models, and on another branch as far as possible from the first one, we glued another set of five larvae corpses of *Operophtera brumata*, an oak-defoliating moth that is often abundant in the study area. The larvae were previously killed by freezing at −80°C. We used glue for securing prey, as it is both the most reliable method for securing corpses, and also the recommended approach to fix plasticine caterpillars by Howe et al. (2009) (36).The prey density was set (i) to reflect the naturally high density of caterpillars at this time of the year, and (ii) to optimize the use of camera traps by ensuring an adequate number of statistical replicates. The five caterpillars on each branch were spaced at least 5 cm apart to remain in the focus of camera traps. Corpses were used instead of live caterpillars for ethical reasons, to avoid gluing live individuals to the branches.

We modeled artificial caterpillars with green plasticine (Staedtler brand, model 8421-5), forming 1.5 × 0.3 cm cylindrical shapes and mimicking Lepidopteran larvae. They had the same color as most plasticine models commonly used in studies of predation interactions in ecology (15–17,23,37) and the same size as *Operophtera brumata* dead larvae used as corpses.

Corpses darken when exposed in the field; after 48 hours, their color differs from that of live individuals. Their smell probably changes too. For these reasons, we exposed fresh models and corpses on a new set of 12 trees every two days. After 16 days, this procedure resulted in a total of 480 model- and 480 corpse- caterpillars installed on 96 oaks.

### Predation analysis

We took plasticine models collected in the field to the laboratory in plastic vials. Note that none of the plasticine models disappeared during the experiment. All 480 models were examined independently by three experienced observers to assess the consistency of their notations (H1). Each observer identified marks of attempted predation by birds using a magnifying glass. The models were considered ‘attacked’ if they exhibited at least one attack mark on the plasticine surface. Observers carefully reread the bite mark guide published by Low et al. (23) beforehand. Following this independent assessment, any cases of doubt regarding an attack was discussed, and observers reached a consensus for testing H2. Hereafter, we refer to the presence of marks reported by human observers’ consensus as ‘predation clues’.

Predation clues on corpses were directly examined in the field, focusing on evidence of bird predation. A missing corpse or one with a missing part was considered a clue of bird attack, following the standard methods in the literature (19,20,24). In contrast, signs such as small holes without significant removal of tissue were attributed to arthropods and were not accounted as bird attack.

### Image analysis

Interactions between predators and corpses as well as models were recorded using camera traps (Brand Ltl Acorn model Ltl-5210M). We installed two camera traps on the trunk of each tree facing the two branches holding models and corpses, respectively. The camera was positioned about 1m from the caterpillars to make it easier to focus (without blurring) (Fig 1). The camera traps were active 24 hours a day for the 14 consecutive days of the experiment. They were set to the highest sensitivity and automatically took bursts of three photos when the passive infrared (PIR) sensors detected motion with a 0.5s trigger speed. At each field visit, we changed the SD cards and checked the batteries to replace them if necessary.

**Fig 1.**
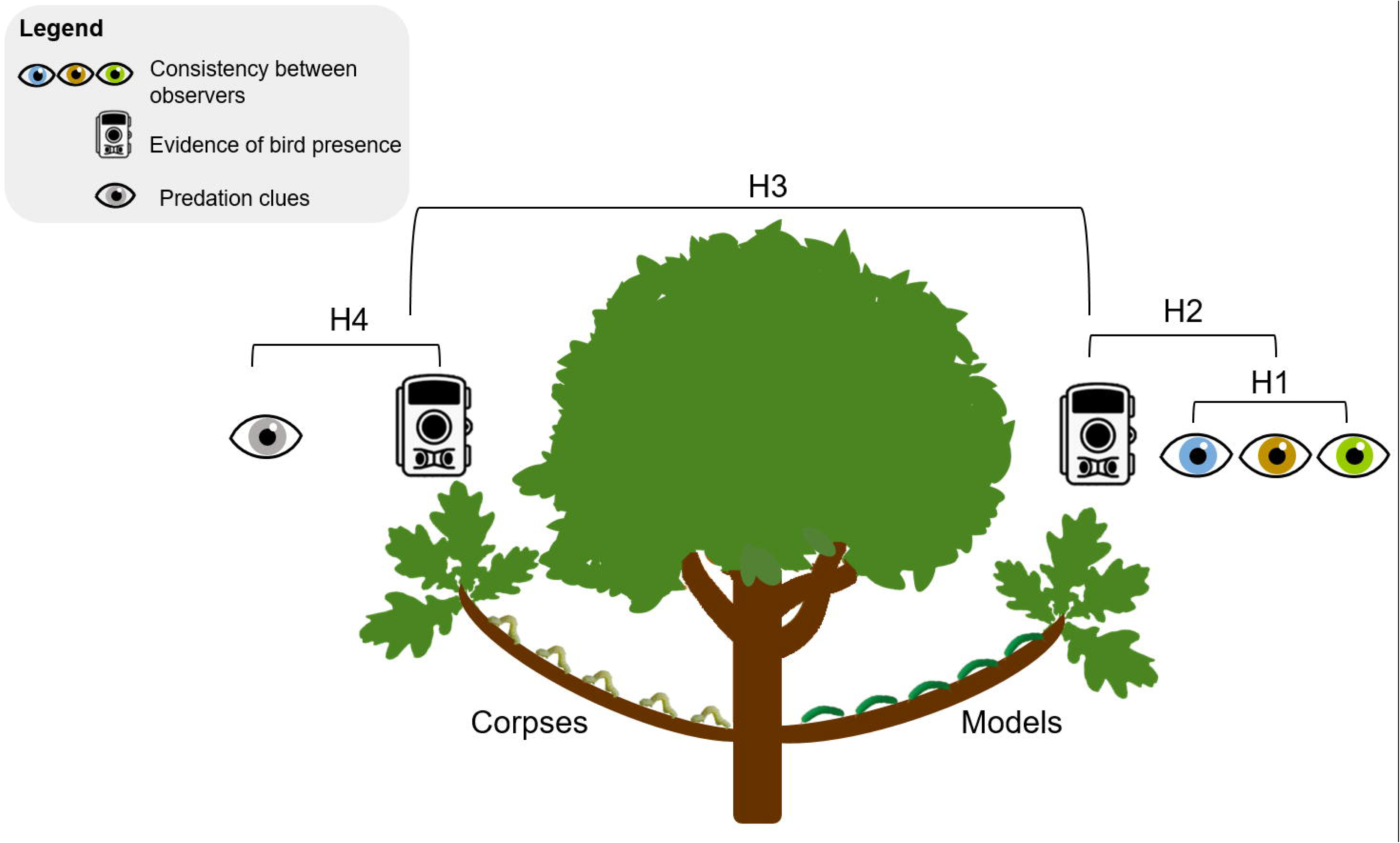
Photographs of the experimental design in the field. Red arrows point toward the camera traps. Blue and yellow arrows point toward five plasticine models (a) and three corpses (b), respectively.

We analyzed the images taken by camera traps using the application EcoAssist (38) supporting the MegaDetector learning model (Machine Learning for Wildlife Image Classification, (39)). This learning model automatically detects the presence of animals in images. We used the most conservative detection confidence threshold of 0.01 for the model in the EcoAssist application, as this threshold previously tested showed a 100% success rate (i.e., all birds present in the images were detected). Following this detection process, we carried out a visual examination by an observer of the images. This examination concerned both corpses and plasticine models.

Our camera traps were able to capture bird presence, including small species (e.g., long-tailed tit), but were insufficient to monitor all predation events due to the birds’ small size and stealthy behavior. Therefore, we defined ‘evidence of predatory bird presence’ at the branch level as the presence of at least one image showing a foraging bird on the branch (hereafter ‘evidence of bird presence’). These images could show individuals in various contexts: birds looking at the camera (Figs 2d and 2e), flying over the target branch (Fig 2a), engaging in foraging activity on the target branch but with natural prey on the bill (Figs 2c and 2e), foraging on experimental prey (Figs 2f and 2h), standing on the experimental branch (Figs 2b and 2g), or feeding during early morning (Fig 2e).

**Fig 2.**
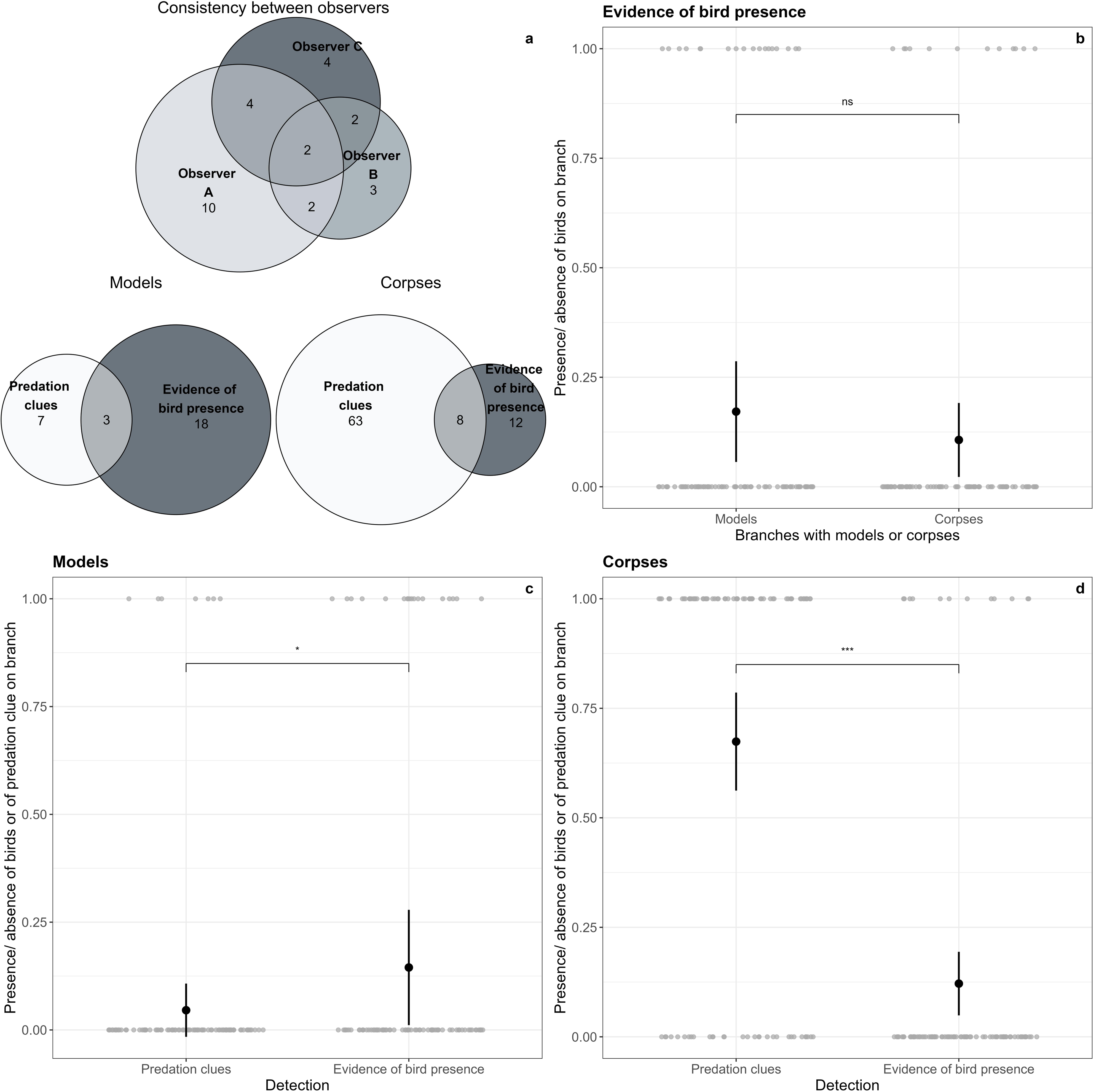
Camera trap photographs of birds taken in various contexts. Birds exhibit different behaviors (perched on a branch, actively foraging, flying, etc., see main text). The blue squares correspond to MegaDetector detections with the associated confidence level. We included red-framed zooms for photos 2b, g, and h, where the birds were difficult to discern. The bird species included are: the Great Tit (*Parus major*, Figs 2a and 2f), the Long-tailed Tit (*Aegithalos caudatus*, Figs 2b and 2g), the European Robin (*Erithacus rubecula*, Fig 2c), the Eurasian Jay (*Garrulus glandarius*, Fig 2d), the European Pied Flycatcher (*Ficedula hypoleuca*, Fig 2e), and the Common Chiffchaff (*Phylloscopus collybita*, Fig 2h)

Subsequently, we statistically compared this evidence of bird presence to the predation clue determined by human observers on the same branch. A branch was categorized as having a predation clue if at least one of its five caterpillars showed a predation clue.

All observed evidence of predator presence, regardless of whether the predator was a bird or a mammal, was also carefully recorded for further discussion. It should be noted, however, that arthropod predators were unlikely to be detected by camera traps. We identified all species associated with the evidence of bird presence.

### Statistical analysis

All analyses were performed in R (40). Fig 3 summarizes all the hypotheses tested using the experimental design.

**Fig 3.**
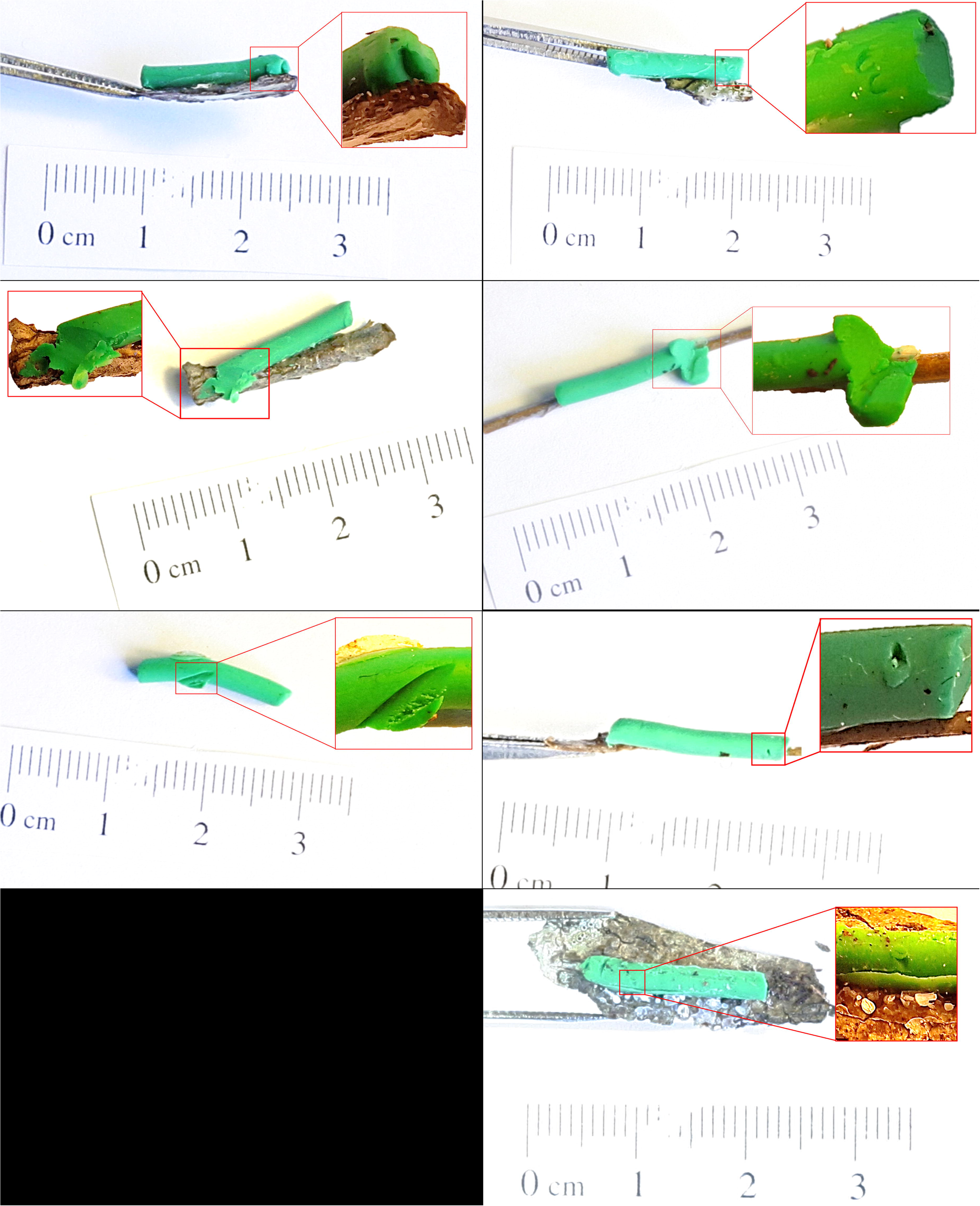
The experimental design testing the four hypotheses. H1: Evaluation of the degree of consistency between observers in the posterior detection of bird predation clues; H2: Comparison between evidence of bird presence assessed by camera traps and predation clues detected by human observers branches with models; H3: Comparison between the evidence of bird presence on corpses versus models both assessed by camera traps; H4: Comparison between evidence of bird presence assessed by camera traps and predation clues detected by human observers branches with corpses.

### Evaluation of the degree of consistency between observers in the posterior detection of bird predation clues (H1)

We relied on the intra-class correlation coefficient (ICC) to assess the degree of consistency between observers in detecting predation clues on plasticine models. The ICC quantifies the degree of agreement between groups or between observers and is bounded between 0 and 1, where 1 indicates perfect intra-group agreement and 0 indicates no agreement beyond what would be expected by chance. According to Koo and Li (41), an ICC value below 0.50 is considered poor, between 0.50 and 0.75 moderate, between 0.75 and 0.90 good, and above 0.90 excellent. We calculated the ICC using the ‘*icc*’ function of the *irr* package (42), with a two-way random effect model because each plasticine model was evaluated by the same set of three observers.

### Comparison between evidence of bird presence assessed by camera traps and predation clues detected by human observers on branches with models and corpses (H2 and H4)

We used two generalized linear mixed models (GLMM, *lme4* package; (43)) with a binomial error distribution to evaluate the bias between observer detection of predation clues and evidence of bird presence assessed by camera traps on (i) branches with models (H2) and on (ii) branches with corpses (H4). In both cases, the detection method (predation clues or evidence of bird presence assessed by a camera) was set as a fixed effect, and tree identity nested in temporal permutation was considered a random factor. The model equation was:

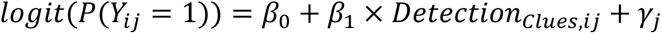

where *Y*_*ij*_ is the binary response variable representing detection at each branch (one data point per detection method and branch). Specifically *Y*_*ij*_ = 1 indicates at least one predation clue or predation evidence on branches with models (H2) or corpses (H4), while *Y*_*ij*_ = 0 indicates no predation clue on any model (H2) or corpse (H4) or no evidence of bird presence. *β*_0_ is the model intercept, representing the logit-transformed probability of detection for camera-assessed bird presence evidence (i.e., *Detection*_*EvidenceBird,ij*_), *β*_1_ is the coefficient of the fixed effect of the predation clues detected by the observer (i.e., *Detection*_*Clues,ij*_), *γ*_*j*_ is the random intercept for tree identity nested in temporal permutation.

### Comparison between the evidence of bird presence on corpses versus models both assessed by camera traps (H3)

We used another GLMM with a binomial distribution to assess the predation bias on models compared to corpses. Fixed effect was the type of prey (model or corpse) and random intercepts were tree identity nested in temporal permutation. The model equation was:

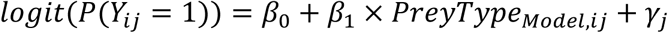

where *Y*_*ij*_ is the binomial response variable indicating bird presence detection at each branch (one data point per type of prey and branch). Specifically, *Y*_*ij*_ = 1 indicates at least one bird presence evidence, while *Y*_*ij*_ = 0 indicates absence of evidence of bird presence. *β*_0_ is the model intercept, representing the logit-transformed probability of detecting bird presence for corpses (i.e., *PreyType*_*Corpse,ij*_), *β*_1_ is the coefficient of the fixed effect of the plasticine models (i.e., *PreyType*_*model,ij*_), *γ*_*j*_ is the random intercept for tree identity nested in temporal permutation.

For both types of GLMMs, we used the ‘*nlminbwrap*’ optimizer, which facilitated model convergence and provided results comparable to the default optimizer of the *lme4* package. It should be noted that since the two GLMMs we used are binomial with a *logit* link function, an inverse transformation of the coefficients must be applied to estimate it as a percentage of predation.

## Results

### Consistency among observers in the detection of bird predation clues (H1)

Observer A detected ten predation clues on 480 plasticine models, whereas Observer B and Observer C detected only three and four predation clues, respectively. At first, the three of them only agreed on two caterpillars (Fig 4a). The intra-class correlation coefficient (ICC) score was 0.47 (CI 95%: 0.41 - 0.52) indicating a poor agreement between the three observers in their assessments of predation clues on the 480 plasticine models (F(479, 960) = 3.61, P < 0.001). The three observers therefore re-assessed predation clues to reach a consensus on the attacked vs non-attacked status. The final, consensual status resulting in seven caterpillars with bird predation clues was used in further analyses.

**Fig 4.**
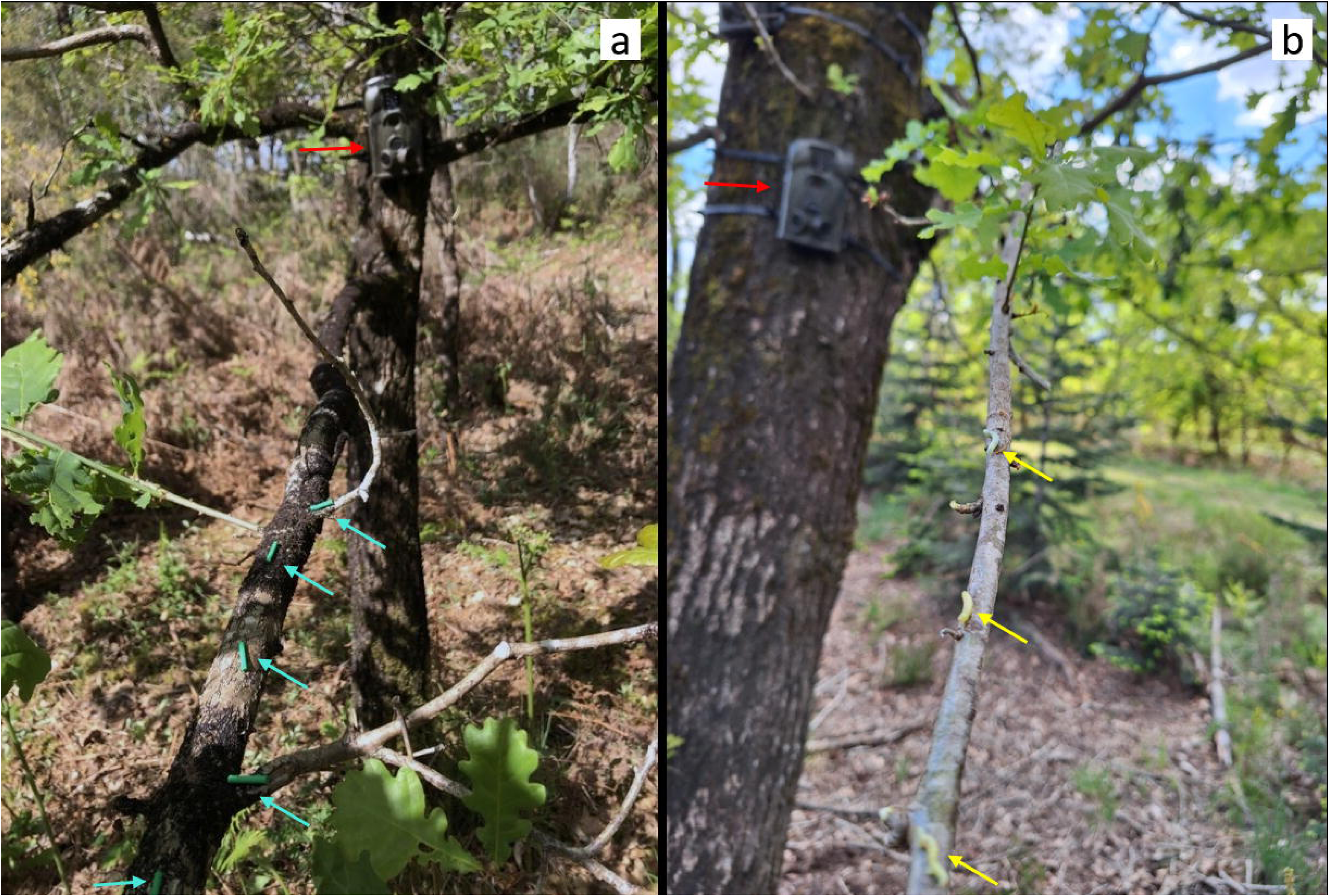
Main results of the study. Euler diagram of consistency between the three observers (a, upper panel), and Euler diagrams of the number of branches with predation clues and with evidence of bird presence for models and corpses (a, lower right and lower left panels, respectively). Results of the generalized mixed-effects models with binomial distribution for the three hypotheses tested: H2 (c), H3 (b), and H4 (d).

### Evidence of bird presence on branches with corpses or models assessed by camera traps (H3)

We obtained 148,157 images from the cameras. The MegaDetector algorithm detected animal presence in 37,495 images (using the lowest detection threshold of 0.01 confidence). After visual evaluation, we ultimately retained only 146 photos showing foraging birds on or around 30 target branches (out of 192). They corresponded to 42 bird individuals belonging to 10 different species (*Aegithalos caudatus*, *Anthus trivialis*, *Phylloscopus collybita*, *Erithacus rubecula*, *Garrulus glandarius*, *Ficedula hypoleuca*, *Cyanistes caerulus*, *Parus major*, *Fringilla coelebs*, and *Turdus philomelos*). We also detected other potential predators on certain branches, such as rodents of the genera *Mus* or *Eliomys* and a Red Squirrel (*Sciurus vulgaris*) (see Figs S1m and z, respectively).

Evidence of bird presence, defined as the presence on images of a foraging bird on or flying toward a branch, did not differ significantly between branches with plasticine models vs. branches with corpses (*β*_1,*Corpse*_ ± SE = −0.5 ± 0.4, *z* = −1.3, *P* = 0.20, Fig 4b). A substantial portion of the variance in bird presence assessment was attributed to random effect (marginal R^2^ = 0.019, conditional R^2^ = 0.19)

### Evidence of bird presence vs. predation clues on branches with models (H2) and corpses (H4)

We found predation clues on 7 plasticine models (7 branches, Fig 5) and 181 corpses (64 branches). On branches with plasticine models, predation clues were associated with bird presence on three branches. We observed the presence of birds on 15 branches with no predation clues on models. On the contrary, predation clues on models were observed on four branches where no birds were detected by camera traps (Fig 4a). Due to missing data caused by camera trap failures on two branches holding corpses and five branches holding models, we excluded these branches from the Euler diagram (Fig 4a), even if predation clues were detected by observers. This explains why, although we identified predation clues on 64 branches holding corpses, the data used to produce the Euler diagram (Fig 4a) were only based on 63 branches.

**Fig 5.**
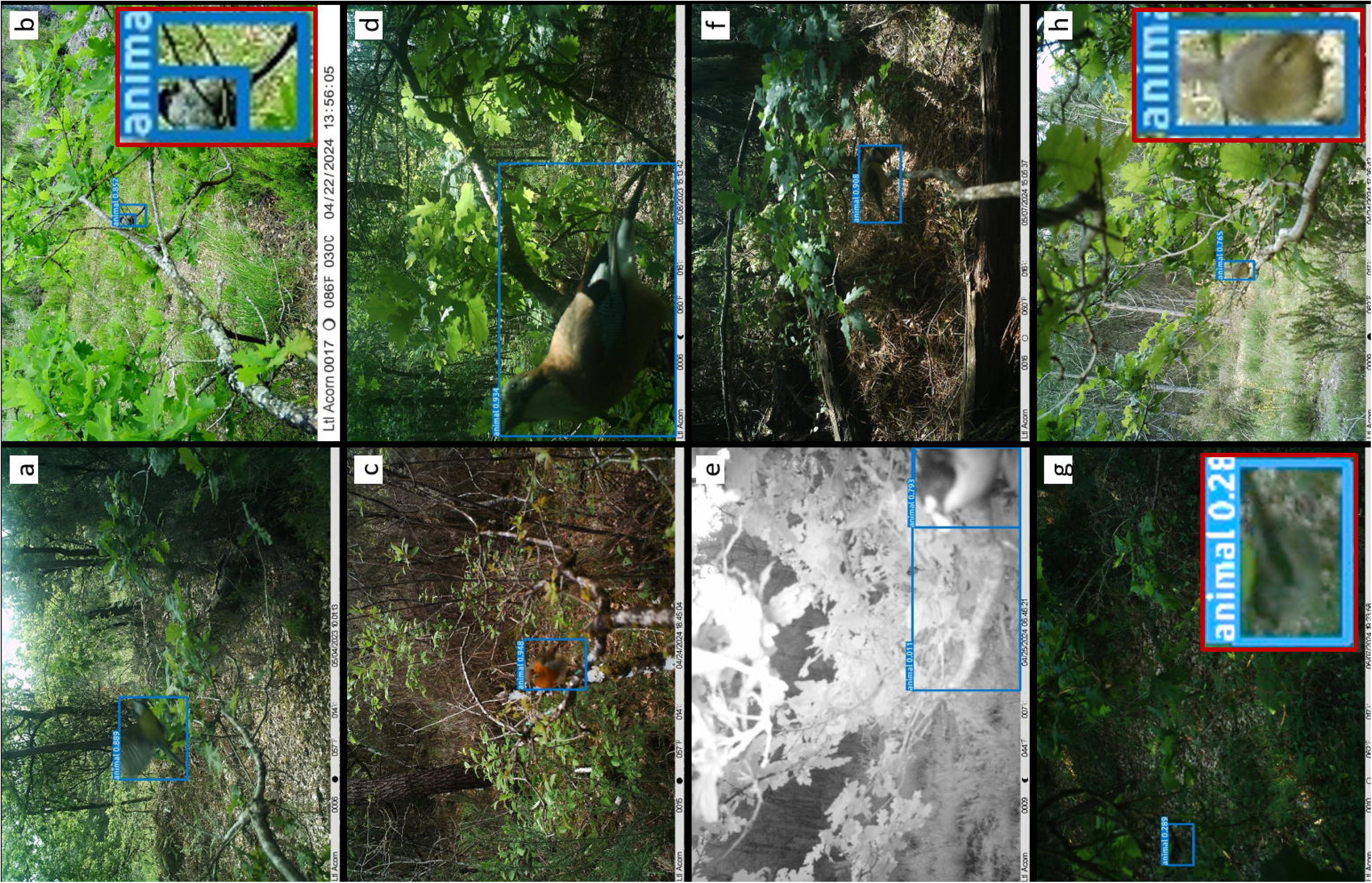
Photographs with mark details. Seven caterpillars that observers considered to have bird predation clues.

For the GLMM analysis, we retained the branches with observed predation clues, even in the absence of corresponding data from camera traps, as GLMMs can account for such imbalances in the data. We detected significantly more evidence of bird presence than predation clues on branches with plasticine models (*β*_1,*Evidence*_ ± SE: 1.3 ± 0.5, *z* = 2.4, *P* < 0.05, Fig 4c). A substantial portion of the variance in predation assessment on branches with plasticine models was attributed to random effect (marginal R^2^ = 0.08, conditional R^2^ = 0.34).

Predation clues on branches with corpses were associated with evidence of bird presence on the same branches in 8 cases. In contrast, we observed evidence of bird presence without predation clues on corpses on 4 branches, and predation clues on corpses without the presence of bird being detected on 56 branches (Fig 4a). There was therefore significantly less evidence of bird presence than predation clues on branches with corpses (*β*_1,*Evidence*_ ± SE: −2.7 ± 0.4, *z* = −6.8, *P* < 0.001, Fig 4d). The random effect accounted for only a small fraction of the variance (marginal R^2^ = 0.35, conditional R^2^ = 0.38).

## Discussion

### Lack of consistency in observations of predation clues on plasticine caterpillar models

Our study confirms the lack of consistency between observers’ evaluation of predation marks on plasticine models. Despite all trained observers re-reading the attack mark guide (23), they did not detect the same number of predation clues. Similar results, even among experienced scientists, were reported by Valdés-Correcher et al. (21) and Castagneyrol et al. (44), who focused specifically on inter-observer consistency with an experimental design suited for that purpose. In our case, the lack of consistency among observers may have been aggravated by the small size of plasticine models mimicking *Operophtera brumata* caterpillars (1.5 × 0.3 cm). These models were smaller than the artificial prey commonly used in the scientific literature (3 cm on average across the five following studies (16,17,19,23,24)), resulting in a perception of predation marks that differs from what we typically observed on larger models. However, this choice was essential for (i) aligning with our target species, which (ii) corresponds to the prey likely to be present locally. We also believe that the size of the artificial prey used in previous studies was sometimes irrelevant when compared to the actual prey sizes present in the study environments. We assume that the insights gained from using realistic model sizes tailored to our study species and context of our study outweigh the limitation of reduced comparability with studies using larger prey models.

To the best of our knowledge, most studies using predation clues on plasticine models rely on a single observer. Information about whether multiple observers could be involved is also rarely provided (e.g., when the primary observer consults one or more colleagues for marks on models that are difficult to assess). Some studies have employed two independent observers evaluating the models by consensus (45,46), which could be a good method to avoid observer bias. It could be also advisable to develop an online standardized training system for identifying bird bill marks on plasticine models, as it is done for insect herbivory (e.g., ZAX herbivory trainer: https://zaxherbivorytrainer.com/(47)). This could complement the comprehensive guide developed by Low et al. (23) and be tailored to different biomes, considering the substantial variations in predator and prey sizes and behaviors.

### Bird presence assessed by camera traps does not differ between branches with plasticine models and corpses

The main result of our study revealed that the evidence of bird presence assessed by camera traps does not significantly differ between experimental branches. Considering that camera traps caught flying birds and birds foraging under poor light conditions (S4 Fig), we assume that the cameras do not miss birds when they are present. Such an outcome would suggest that plasticine models can be used to estimate bird predation, as they offer the additional advantage of overcoming the ethical issues associated with the use of real prey, whether dead or alive. This result differs from our findings regarding the number of predation clues assessed by observers and contradicts other studies that find more bird attacks on real prey compared to plasticine ones (19,20).

Specifically, we observed a ratio of 7 predation clues on plasticine models to 181 on corpses. Initially, we did not consider arthropods capable of removing an entire corpse or significant portions of it. However, based on our results, we are compelled to consider this possibility. Similar to the findings by Zvereva & Kozlov (20), we believe that the corpses could have been removed and attacked by arthropods as we found ants crawling around 24 corpses out of 480, but other predators such as spiders or other arthropods could also have been involved. It is also possible that the corpses attracted necrophages, although we have no formal evidence of this. These findings are also in accordance with Nimalrathna et al. (24) who found that the attack rate by vertebrate predators was lower on dead prey than on live prey and plasticine models, while invertebrates predated most on dead prey followed by live prey and plasticine models. Considering our and Nimalrathna et al. (24) results, we conclude that our observations of predation on corpses could have been driven by arthropod consumers.

Although it may be tempting to conclude that birds do not distinguish between real and plasticine caterpillars, and thus claim this method as reliable, it must be noted that we cannot rely on (i) the posterior evaluation of clues on corpses (plasticine models probably allow for a clearer distinction between bird marks and other types of marks compared to real prey); and (ii) the evidence of bird presence assessed by camera traps as definitive evidence of predation. Due to these remaining uncertainties, it seems particularly important to be able to effectively evaluate the plasticine model method, which is currently limited by available technology. To our knowledge, there is currently no reliable, affordable technique for accurately identifying birds preying on insects. Camera traps available on the market have motion detection speeds that are too slow for birds or do not allow for fast sequences of pictures or continuous recording. Cameras that do offer continuous recording have other limitations in data storage, power supply, and encoding time. There is thus a need for advanced cameras that allow us to observe fast behavioral phenomena of relatively small animals, such as foraging birds. These cameras could be used in various fields such as trophic interaction ecology, conservation ecology, and behavioral ecology. Although some initiatives offering prototypes exist (e.g., (48)), the costs are currently still too high to be affordable for scientific purposes.

The most limiting trade-off is between the camera performance and energy consumption. For scientific research, we need at least 24 hours of endurance, which is currently not possible for continuous recording. The cameras could therefore operate by capturing a sequence of pictures at intervals of at least 0.5 seconds continuously to maximize the chance of capturing the moment of predation (86,400 pictures/day). At the same time, the batteries must be removable for quick changes in the field. Additionally, the cameras should be affordable, relatively small, light, and waterproof, and finally have programmed a short encoding time.

### Predation clues assessed by observers do not correlate with bird presence assessed by camera traps

Predation clues assessed by observers on plasticine models or corpses did not reflect bird foraging activity as revealed by camera traps. Another study comparing camera traps and posterior detection to assess predation on artificial snakes also revealed that predatory birds were underrepresented in the videos triggered after motion detection compared to the number of predation marks on artificial replicates (49). Moreover, O’Brien & Kinnaird (50) demonstrated in 2008 that camera traps were more suitable for larger birds such as pheasants than for functionally insectivorous birds targeted in our study. We acknowledge that modern camera trap technology has improved considerably since 2008, although we are not aware of any recent studies that have specifically addressed this issue in the context of bird predation.

Contrary to our hypothesis, we found more evidence of bird presence than predation clues on plasticine models suggesting that fake prey underestimates the foraging activity of avian insectivores. Considering the particularly high density of live prey during this time of year (L.S., N.P., A.M. pers. observation), an explanation could be that experienced birds prefer live caterpillars found on the same branch over the models (see also (20)). It is, however, important to consider the low numbers of both observations (clues vs. evidence) (Fig 4c) implying a weak indicative value. Moreover, not all evidence of bird presence may necessarily reflect predation attempts.

The glue used to attach the prey sometimes caused such small models to harden, potentially preventing the later detection of predation clues. Finally, we could not exclude curiosity-driven behaviors from some individuals, notably Eurasian jays, who seemed to gaze intently at the camera rather than at the available prey. While we anticipated this curiosity behavior, we expected it to manifest more in predation clues on the models (e.g., such as claw marks). We carefully re-checked models on which no predation clues were initially identified, whereas we obtained evidence of birds foraging on the corresponding branches. The three observers confirmed no predation clue was visible on these models.

We observed more predation clues (181) than presence evidence on corpses (12). As described above, we believe that the effect size is likely driven by arthropod consumers. This assumption, however, cannot be currently tested as we realized that we are probably not able to distinguish between arthropod and bird clues on corpses, and at the same time we cannot rely on the evidence of bird presence to be evidence of a genuine predation event. Additionally, we note that photographs from the camera traps revealed potentially predatory rodents on one of the branches with corpses.

### Perspectives

Overall, the results discussed here provide only a limited window to the interpretation of whether the use of plasticine models should be regarded as a reliable method. Therefore, we encourage future protocols to explore other methods, such as considering behavioral studies in controlled conditions offering multiple types of prey to birds. Still, our study paves the way for the use of new technologies, such as passive monitoring methods (e.g., video surveillance), to obtain real evidence of predation. We, therefore, advocate for their development and use, which could eventually refine the widely used plasticine model method in the ecology of trophic interactions.

It is important to note that in this study, camera traps were used as a way to evaluate the plasticine model method rather than as a tool to directly assess predation. In the context of future methodological developments, however, cameras could potentially be used to assess predation when combined with plasticine models or corpses. Taking into account all results and technical difficulties of this study as well as the results of other studies, we summarized the advantages and disadvantages of each method, in their current state of development, for quantifying bird predation, along with future recommendations in Table 1.

**Table 1.** Advantages, disadvantages, comments, and recommendations for the three possible methods of assessing bird predation: plasticine models, real prey, and cameras in their actual state of development. Note that cameras must be used in combination with one of the other two methods. If a cell in the third column spans two rows in the first column labeled ‘Methods,’ it applies to both methods. Literature references supporting our arguments are marked in blue.

| Methods | Advantages | Disadvantages | Comments/ recommendations |
| --- | --- | --- | --- |
| ***Plasticine models*** | - Low-cost<br>- Easy method<br>- Standardized<br>- Ethical<br>- Posterior detection possible<br>- Bird predation marks are relatively distinct from arthropod marks | - Artificiality: immobile, too bright color resulting in unnatural contrast with branch; non-natural smell; often oversized regarding the size of the species naturally occurring in the habitat<br>- Inconsistency of human detection | - Most models used are too large compared to native European caterpillars ((14); see S5 Fig). Therefore, we recommend adjusting the models' size according to the species likely to be present at study sites.<br>- The vivid green color is not always the most suitable choice for plasticine. We recommend adjusting the color of larvae in a given geography and habitat (e.g., (51)).<br>- We recommend several independent observers (e.g., (45,46)) or an online training system (on the model of (47)) to mitigate the observer detection bias. |
|  |  |  | - It is necessary to consider a realistic density of models concerning the actual abundance of prey in the study area at any given time (14).<br>- Consider exposing the prey (models or real caterpillars) according to the natural behavior of the species they represent (e.g., petioles of<br>the leaves, inner space of the rolled leaves, tree trunks, etc.) (e.g., (15,17,27)).<br>- Consider the appropriate attachment method regarding the size of the prey, (e.g., the glue may not be the best solution in the case of small prey and could be substituted with pins or double-sided adhesive tape) (e.g., (15,17,27,52)). |
| **Real prey** | - Really reflects predation | - Expensive and time-consuming rearing<br>- Unethical<br>In the case of using dead natural prey :<br>- Corpses must be changed frequently<br>- The predation marks of different taxa are indistinguishable<br>- Corpses may attract necrophages | - Use live prey to better reflect predation. Develop protocols to utilize live prey without attachment for ethical reasons while preventing their escape from branches (e.g., (19,20,53)). |
|  |  |  | - Combination of real prey and cameras. |
| **Camera traps** | - Theoretically allows obtaining evidence of predation by birds (but this study showed that current camera traps do not yet allow this)<br>- Identification of bird species<br>- Detection of other vertebrate predators<br>- Knowing the exact time of predation | - Expensive (54)<br><u>Currently:</u><br>- Passive infrared sensors are not suitable for small birds (50)<br>- Too slow motion detection speeds for photos and even slower for videos<br>- Only a few photos in burst mode after motion detection decrease the chance of capturing the predation event<br>- Too short video times<br>- Non-existing long-term continuous recording<br>- Low data storage<br>- Short-term power supply | - We strongly encourage programmers and electronics engineers to use low-cost microcontrollers coupled with small cameras and high-capacity batteries to obtain real evidence of not only bird predation but generally animal behavior under natural conditions. Such a monitoring method could allow us to conduct reproducible experiments on a large geographic scale, in different habitats, and with multiple target larva species to model an overall bias of the plasticine model method. |
|  |  | - Long encoding time<br>- High power supply (54)<br>- Need for artificial intelligence to process thousands of photos (48) |  |

## Conclusion

Our study sheds new light on the evaluation of the plasticine model method for characterizing bird predation. It suggests that predation clues do not accurately reflect the presence of predatory birds as assessed by camera traps, on either the branches bearing plasticine models or those bearing dead natural caterpillars. As we found no differences in the behavioral responses of birds to artificial prey compared with dead natural ones, we would conclude that using caterpillar corpses did not present advantages over using plasticine models for assessing avian predation on folivores. However, the inter-observer inconsistencies in assessing predation clues on plasticine models, implying an extent of subjectivity, prevent us from claiming this method is fully reliable. Existing cameras do not yet allow to evaluate the plasticine caterpillar method. Without advanced technologies to assess the biases inherent in using plasticine models and to accurately measure predation, we argue that plasticine models offer a simpler, more economical, ethical, and equally reliable alternative to real larvae for studying bird predation. Therefore, we also advocate for the development of more precise monitoring devices and more global initiatives building upon our study, with the aim of improving these methods once reliable and effective monitoring tools become available.

## Acknowledgments

We thank Alba Lazaro-Gonzalez, Heidy Schimann, Inge Van Halder, Tom Barlier, Thomas Ribot, and Thibaud Coupart for their help in the field. We also thank to Pavel Potocký, who collected fertilized autumnal moth females.

## Data availability plan

The R Markdown format will be used to ensure that the statistical analyses are fully reproducible, with the code and data made publicly available (https://recherche.data.gouv.fr/) upon acceptance of the study, following the FAIR principles for scientific data management.

## Supporting Information

**S1 Table. Experimental design of the manuscript “Accuracy in bird predation assessment: Camera traps testing the efficacy of plasticine caterpillars as prey models” as described in the registered report prior to implementation of the experiment.**

It should be noted that when drafting the protocol for the registered report, we initially thought we would be able to detect predation evidence with camera traps at the caterpillar level. Later, we realized that this evidence was only detectable at the branch level, which led to slight adjustments in the statistical models in the final manuscript. See the Materials and Methods section for further details.

**S2 Fig. The results of power analyses with 1000 simulations for the three different scenarios tested (hypotheses 2 and 4).**

The power analysis results according to the three different datasets are the following: predation clues 5% - 10% predation evidence: 99.80% (range: 99.28, 99.98); clues 8% - 9% evidence: 99.90% (range: 99.44, 100.00); clues 9% - 9% evidence: 0.00% (range: 0.00, 0.37).

**S3 Fig. The results of power analyses with 1000 simulations for the 3 different scenarios tested (hypothesis 3).**

The power analyses were based on the sample size of 420 caterpillars per fixed factor. The power analysis results according to the three different datasets are the following: model 5% - 10% corpse: 83.30% (range: 80.84, 85.56); model 5% - 20% corpse: 100.00% (range: 99.63, 100.00); model 10% - 20% corpse: 99.30% (range: 98.56, 99.72).

**S4 Fig. All detected birds by MegaDetector AI software.**

Camera traps photographs of potential predators of caterpillars. The blue squares correspond to MegaDetector detections with the associated confidence level. The animal species included are: the Long-tailed Tit (*Aegithalos caudatus*, S1a and S1ag Figs), the Tree Pipit (*Anthus trivialis*, S1c and S1k Figs), the Common Chiffchaff (*Phylloscopus collybita*, S1b and S1g Figs), the European Robin (*Erithacus rubecula*, S1d, S1f, S1j, S1o, S1p, S1q, S1s, S1ab, and S1ad Figs), the Eurasian Jay (*Garrulus glandarius*, S1e, S1l, S1r, S1y, S1af, S1ak, S1an, and S1ap Figs), the European Pied Flycatcher (*Ficedula hypoleuca*, S1h and S1i Fig.), the Red Squirrel (*Sciurus vulgaris*, S1m Fig), the Eurasian Blue Tit (*Cyanistes caerulus*, S1n and S1am Figs), the Great Tit (*Parus major*, S1t, S1v, S1w, S1aa, S1ac, S1ah and S1ai Figs), the Common Chaffinch (*Fringilla coelebs*, S1u, S1x, S1ae, S1aj, S1al, S1ao and S1aq Figs), unidentified rodents (S1z Fig), and the Song Trush (*Turdus philomelos*, S1ar Fig).

**S5 Fig. Photographs of larvae found in the study area.**

From top to bottom, the families, genera, or species are: *Cydalima perspectalis* (non-native species), Geometridae family, Tenthredinidae family (likely of Caliroa genus), Tenthredinidae family (likely of Periclista genus), *Operopthera brumata*. Scale in cm.

## References

1. Díaz-Siefer P, Olmos-Moya N, Fontúrbel FE, Lavandero B, Pozo RA, Celis-Diez JL. Bird-mediated effects of pest control services on crop productivity: a global synthesis. J Pest Sci. 2022 Mar;95(2):567–76.

2. Remmel T, Davison J, Tammaru T. Quantifying predation on folivorous insect larvae: the perspective of life-history evolution. Biol J Linn Soc. 2011 Sep;104(1):1–18.

3. Ford AT, Goheen JR. Trophic Cascades by Large Carnivores: A Case for Strong Inference and Mechanism. Trends Ecol Evol. 2015 Dec;30(12):725–35.

4. Nyffeler M, Şekercioğlu ÇH, Whelan CJ. Insectivorous birds consume an estimated 400– 500 million tons of prey annually. Sci Nat. 2018 Aug;105(7–8):47.

5. Sherry TW, Kent CM, Sánchez NV, Şekercioğlu ÇH. Insectivorous birds in the Neotropics: Ecological radiations, specialization, and coexistence in species-rich communities. The Auk. 2020 Dec 24;137(4):ukaa049.

6. Thiollay JM. Comparative Foraging Success of Insectivorous Birds in Tropical and Temperate Forests: Ecological Implications. Oikos. 1988 Jul;53(1):17.

7. Collins CT, Watson A. Field Observations of Bird Predation on Neotropical Moths. Biotropica. 1983 Mar;15(1):53.

8. Mäntylä E, Kleier S, Lindstedt C, Kipper S, Hilker M. Insectivorous Birds Are Attracted by Plant Traits Induced by Insect Egg Deposition. J Chem Ecol. 2018 Dec;44(12):1127–38.

9. Poloni R, Dhennin M, Mappes J, Joron M, Nokelainen O. Exploring polymorphism in a palatable prey: predation risk and frequency dependence in relation to distinct levels of conspicuousness. Evol Lett. 2024 Jan 11;qrad071.

10. van Bael SA, Philpott SM, Greenberg R, Bichier P, Barber NA, Mooney KA, et al. Birds as predators in tropical agroforestry systems. Ecology. 2008 Apr;89(4):928–34.

11. Michalski M, Nadolski J, Marciniak B, Loga B, Bańbura J. Faecal Analysis as a Method of Nestling Diet Determination in Insectivorous Birds: A Case Study in Blue Tits *Cyanistes caeruleus* and Great Tits *Parus major*. Acta Ornithol. 2011 Dec;46(2):164–72.

12. Sam K, Koane B, Jeppy S, Sykorova J, Novotny V. Diet of land birds along an elevational gradient in Papua New Guinea. Sci Rep. 2017 Mar 9;7(1):44018.

13. Brodie III ED. Differential avoidance of coral snake banded patterns by free-ranging avian predators in Costa Rica. Evolution. 1993;47(1):227–35.

14. Lövei GL, Ferrante M. A review of the sentinel prey method as a way of quantifying invertebrate predation under field conditions: Measuring predation pressure by sentinel prey. Insect Sci. 2017 Aug;24(4):528–42.

15. Roslin T, Hardwick B, Novotny V, Petry WK, Andrew NR, Asmus A, et al. Higher predation risk for insect prey at low latitudes and elevations. Science. 2017 May 19;356(6339):742–4.

16. Schillé L, Valdés-Correcher E, Archaux F, Bălăcenoiu F, Bjørn MC, Bogdziewicz M, et al. Decomposing drivers in avian insectivory: Large-scale effects of climate, habitat and bird diversity. J Biogeogr. 2024 Feb 4;jbi.14808.

17. Valdés-Correcher E, Moreira X, Augusto L, Barbaro L, Bouget C, Bouriaud O, et al. Search for top-down and bottom-up drivers of latitudinal trends in insect herbivory in oak trees in Europe. Keith S, editor. Glob Ecol Biogeogr. 2021 Mar;30(3):651–65.

18. Bateman PW, Fleming PA, Wolfe AK. A different kind of ecological modelling: the use of clay model organisms to explore predator–prey interactions in vertebrates. J Zool. 2017 Apr;301(4):251–62.

19. Rodriguez-Campbell A, Rahn O, Chiuffo MC, Hargreaves AL. Clay larvae do not accurately measure biogeographic patterns in predation. J Biogeogr. 2023;

20. Zvereva EL, Kozlov MV. Predation risk estimated on live and artificial insect prey follows different patterns. Ecology. 2023 Mar;104(3):e3943.

21. Valdés-Correcher E, Mäntylä E, Barbaro L, Damestoy T, Sam K, Castagneyrol B. Following the track: accuracy and reproducibility of predation assessment on artificial caterpillars. Entomol Exp Appl. 2022;170(10):914–21.

22. Rößler DC, Pröhl H, Lötters S. The future of clay model studies. BMC Zool. 2018 Dec;3(1):6.

23. Low PA, Sam K, McArthur C, Posa MRC, Hochuli DF. Determining predator identity from attack marks left in model caterpillars: guidelines for best practice. Entomol Exp Appl. 2014 Aug;152(2):120–6.

24. Nimalrathna TS, Solina ID, Mon AM, Pomoim N, Bhadra S, Zvereva EL, et al. Estimating predation pressure in ecological studies: controlling bias imposed by using sentinel plasticine prey. Entomol Exp Appl. 2023 Jan;171(1):56–67.

25. Valdés-Correcher E, van Halder I, Barbaro L, Castagneyrol B, Hampe A. Insect herbivory and avian insectivory in novel native oak forests: Divergent effects of stand size and connectivity. For Ecol Manag. 2019 Aug;445:146–53.

26. Muchula K, Xie G, Gurr GM. Ambient temperature affects the utility of plasticine caterpillar models as a tool to measure activity of predators across latitudinal and elevational gradients. Biol Control. 2019 Feb;129:12–7.

27. Mrazova A, Houska Tahadlová M, Řehová V, Sam K. The specificity of induced chemical defence of two oak species affects differently arthropod herbivores and arthropod and bird predation. Arthropod-Plant Interact. 2023 Apr;17(2):141–55.

28. Campos RC, Steiner J, Zillikens A. Bird and mammal frugivores of *Euterpe edulis* at Santa Catarina island monitored by camera traps. Stud Neotropical Fauna Environ. 2012 Aug;47(2):105–10.

29. Zhu C, Li W, Gregory T, Wang D, Ren P, Zeng D, et al. Arboreal camera trapping: a reliable tool to monitor plant-frugivore interactions in the trees on large scales. Remote Sens Ecol Conserv. 2022;8(1):92–104.

30. Schillé L, Plat N, Barbaro L, Jactel H, Raspail F, Rivoal JB, et al. Accuracy in bird predation assessment: Camera trap testing the efficacy of plasticine caterpillars as prey models. PCI RR Manag Board [Internet]. 2024 Jun 12 [cited 2024 Nov 14]; Available from: https://osf.io/https://osf.io/be2hn

31. Schille L, Valdés-Correcher E, Archaux F, Bălăcenoiu F, Bjørn MC, Bogdziewicz M, et al. Data and codes for the article “Decomposing drivers in avian insectivory: large-scale effects of climate, habitat and bird diversity” [Internet]. Recherche Data Gouv; 2024 [cited 2024 Feb 2]. Available from: https://entrepot.recherche.data.gouv.fr/citation?persistentId=doi:10.57745/0E0JEA

32. Naef-Daenzer B. Radiotracking of great and blue tits: new tools to assess territoriality, home-range use and resource distribution. 1994;

33. Lerche-Jørgensen M, Mallord JW, Willemoes M, Orsman CJ, Roberts JT, Skeen RQ, et al. Spatial behavior and habitat use in widely separated breeding and wintering distributions across three species of long-distance migrant *Phylloscopus* warblers. Ecol Evol. 2019 Jun;9(11):6492–500.

34. Cornell Laboratory of Ornithology. Birds of the World. In: Billerman SM, Keeney BK, Rodewald PG, Schulenberg TS, editors. Cornell Laboratory of Ornithology, Ithaca, NY, USA [Internet]. 2022. Available from: https://birdsoftheworld.org/bow/home

35. Naguib M, Titulaer M, Waas JR, Van Oers K, Sprau P, Snijders L. Prior territorial responses and home range size predict territory defense in radio-tagged great tits. Behav Ecol Sociobiol. 2022 Mar;76(3):35.

36. Howe A, Lövei GL, Nachman G. Dummy caterpillars as a simple method to assess predation rates on invertebrates in a tropical agroecosystem. Entomol Exp Appl. 2009 Jun;131(3):325–9.

37. Sam K, Koane B, Novotny V. Herbivore damage increases avian and ant predation of caterpillars on trees along a complete elevational forest gradient in Papua New Guinea. Ecography. 2015;38(3):293–300.

38. van Lunteren P. EcoAssist: A no-code platform to train and deploy custom YOLOv5 object detection models. J Open Source Softw. 2023;8(88):5581.

39. Beery S, Morris D, Yang S. Efficient Pipeline for Camera Trap Image Review [Internet]. arXiv; 2019 [cited 2024 Mar 16]. Available from: http://arxiv.org/abs/1907.06772

40. R Core Team. R: A Language and environment for statistical computing. [Internet]. R Foundation for Statistical Computing, Vienna, Austria; 2021. Available from: https://www.r-project.org/

41. Koo TK, Li MY. A Guideline of Selecting and Reporting Intraclass Correlation Coefficients for Reliability Research. J Chiropr Med. 2016 Jun;15(2):155–63.

42. Gamer M, Lemon J, Singh IFP. irr: Various Coefficients of Interrater Reliability and Agreement [Internet]. 2019 [cited 2024 Mar 17]. Available from: https://cran.r-project.org/web/packages/irr/index.html

43. Bates D, Mächler M, Bolker B, Walker S. Fitting Linear Mixed-Effects Models Using **lme4**. J Stat Softw. 2015;67(1):5–8.

44. Castagneyrol B, Valdés-Correcher E, Bourdin A, Barbaro L, Bouriaud O, Branco M, et al. Can School Children Support Ecological Research? Lessons from the *Oak Bodyguard* Citizen Science Project. Citiz Sci Theory Pract. 2020 Mar 18;5(1):10.

45. Czarnecki C, Manderino R, Parry D. Reduced avian predation on an ultraviolet-fluorescing caterpillar model. Can Entomol. 2022;154(1):e10.

46. Franco JC, Branco M, Conde S, Garcia A, Fernandes MR, Lima Santos J, et al. Ecological Infrastructures May Enhance Lepidopterans Predation in Irrigated Mediterranean Farmland, Depending on Their Typology and the Predator Guild. Sustainability. 2022 Mar 25;14(7):3874.

47. Xirocostas ZA, Debono SA, Slavich E, Moles AT. The ZAX Herbivory Trainer—Free software for training researchers to visually estimate leaf damage. Methods Ecol Evol. 2022;13(3):596–602.

48. Darras KFA, Balle M, Xu W, Yan Y, Zakka VG, Toledo-Hernández M, et al. Eyes on nature: Embedded vision cameras for terrestrial biodiversity monitoring. Methods Ecol Evol. 2024 Oct 28;2041-210X.14436.

49. Akcali CK, Adán Pérez-Mendoza H, Salazar-Valenzuela D, Kikuchi DW, Guayasamin JM, Pfennig DW. Evaluating the utility of camera traps in field studies of predation. PeerJ. 2019 Feb 25;7:e6487.

50. O’Brien TG, Kinnaird MF. A picture is worth a thousand words: the application of camera trapping to the study of birds. Bird Conserv Int. 2008 Sep;18(S1):S144–62.

51. Zvereva EL, Castagneyrol B, Cornelissen T, Forsman A, Hernández-Agüero JA, Klemola T, et al. Opposite latitudinal patterns for bird and arthropod predation revealed in experiments with differently colored artificial prey. Ecol Evol. 2019 Dec;9(24):14273– 85.

52. Richards LA, Coley PD. Seasonal and habitat differences affect the impact of food and predation on herbivores: a comparison between gaps and understory of a tropical forest. Oikos. 2007 Jan;116(1):31–40.

53. Sam K, Jorge LR, Koane B, Amick PK, Sivault E. Vertebrates, but not ants, protect rainforest from herbivorous insects across elevations in Papua New Guinea. J Biogeogr. 2023 Jul 19;jbi.14686.

54. Glover-Kapfer P, Soto-Navarro CA, Wearn OR. Camera-trapping version 3.0: current constraints and future priorities for development. Remote Sens Ecol Conserv. 2019;5(3):209–23.

