## Supplemental Table 1, S2 Fig, S3 Fig, S4 Fig and S5 Fig for "Camera traps unable to determine whether plasticine models of caterpillars reliably measure bird predation"

### Supporting Information

**S1 Table. Experimental design of the manuscript "Accuracy in bird predation assessment: Camera traps testing the efficacy of plasticine caterpillars as prey models" as described in the registered report prior to implementation of the experiment.**
It should be noted that when drafting the protocol for the registered report, we initially thought we would be able to detect predation evidence with camera traps at the caterpillar level. Later, we realized that this evidence was only detectable at the branch level, which led to slight adjustments in the statistical models in the final manuscript. See the Materials and Methods section for further details.

| **Question** | **Hypothesis** | **Sampling plan** | **Analysis Plan** | **Rationale for deciding the sensitivity of the test for confirming or disconfirming the hypothesis** | **Interpretation given different outcomes** | **Theory that could be shown wrong by the outcomes** |
|---|---|---|---|---|---|---|
| 1. Could the lack of consistency between observers be detrimental to the plasticine models method for assessing bird predation? | H1. The experienced observers who underwent the same training show a high degree of consistency in assessing attacks on models. | The 3 observers will carefully read the plasticine models bite image guide by Low et al. 2014 beforehand.<br><br>Each of the 3 observers will then evaluate each of the 420 models and define whether or not it has been attacked by a bird. | The consistency of the 3 observers' assessments will be calculated using the intra-class correlation coefficient (ICC) with a two-way random effect model because each model will be evaluated by the same set of 3 observers. | We will use the *icc* function of the "irr" package (Gamer et al., 2019) to calculate the ICC among observers. We will consider an ICC greater than 0.75 to be representative of good consistency between the 3 observers (Koo & Li, 2016). In this case, we argue that we can randomly choose one observer among the 3. Its evaluation will be called "predation clues" for the following hypothesis. | If the ICC between the 3 observers is less than 0.75, their consistency will be considered insufficient (Koo & Li, 2016). In such a case, for each model where there was doubt as to its attacked or non-attacked status, the 3 observers will discuss and agree on its status. We believe that scientists using plasticine models probably proceed in this way, seeking the advice of colleagues when in doubt. | Theoretically, the plasticine model method is based on very good consistency between observers, without which human judgment would be too unreliable and subjective to characterize predation interactions using this method. Should this be the case, we would discuss the need for a standardized training system for recognizing bird bill marks on models. |

| | | | | | | |
|---|---|---|---|---|---|---|
| 2. Is there a bias between observer and camera detection of bird marks on models? | H2. Posterior detection of marks on models is inaccurate, increasing the number of attacks, because observers unconsciously account for marks not related to predation. | We will select 12 pedunculate oaks. We will attach 5 plasticinemodels to one of the branches of each of them. We will attach 5 corpses (dead caterpillars) on another branch of each of them, as far as possible from the first branch. A camera trap will face each branch bearing a set of 5 caterpillars, whether corpses or models. Every two days, the caterpillars will be collected and a new set of caterpillars will be installed on 12 new oaks (this process will be referred to hereafter as "permutation").<br><br>(H2) Bird predation evidence will be recorded on each of | $logit(P(Y_{ijk} = 1)) = \beta_0 + \beta_1 \times Detection_{Clues,ijk} + \gamma_j + \zeta_k$<br><br>where $Y_{ijk}$ is the binary response variable (one data point per detection method, branch, and model), with $Y_{ijk}=1$ indicating at least one predation clue or one bird presence evidence, and $Y_{ijk}=0$ indicating no detection. $\beta_0$ is the model intercept, representing the logit-transformed probability of detection for the camera-assessed bird presence evidence (i.e., $Detection_{Evidence,ijk}$), $\beta_1$ the coefficient of the fixed effect predation clues detected by the observer (i.e., $Detection_{Clues,ijk}$). $\gamma_j$ is the random effect for tree identity nested in temporal permutation and follows a normal law $\gamma_j \sim N(0,\sigma_\gamma^2)$, $\zeta_k$ is the random effect for model identity and follows a normal law $\zeta_k \sim N(0,\sigma_\zeta^2)$. | In the experimental area, a previous study using plasticine caterpillar models coupled with observer detection of predation clues found bird attack rates ranging from 5 to 10% (see data from site "ele.val_1", Schille et al., 2024).<br><br>We used these references in conjunction with our hypothesis and sample size to simulate 3 different datasets with different effect sizes:<br><br>(1) Predation evidence of 5% detected with the camera versus 10% predation clues detected by human; (2) 8% evidence vs. 9% clues; (3) 9% evidence vs. 9% clues.<br><br>Note that the simulation of this dataset is not fully randomized. We simulated the attacked/non-attacked status of the vast majority of models identical between the two types of detection. Only 1% of | If $\beta_1 \simeq 0$, predation clues reflect predation evidence assessed by the camera because the difference between predation clues and evidence would then be minimal, and $\beta_1$ would not be significant.<br><br>If $\beta_1 < 0$, predation clues are underestimated in relation to predation evidence assessed by the cameras. | If predation clues overestimate or underestimate real predation on plasticine models, observers should (1) go through a process of training and/or assessment of their ability to detect predation marks and/or (2) consider the more systematized use of cameras to observe what is actually happening in the field. |

| | | | | |
|---|---|---|---|---|
| | | the 420 models using camera traps. In parallel, one or more observers (depending on the consistency of their assessments, see previous hypothesis) will also have assessed bird predation clues on each caterpillar. Each model will therefore be considered attacked or not attacked by 2 types of detection (clues or evidence). | We predict $\beta_1 > 0$ indicating an increased detection rate of predation clues compared to bird presence evidence.<br><br>The mathematical transformation<br><br>$p = \frac{\exp(\beta_0 + \beta_1 \times Detection_{ijk})}{1+\exp(\beta_0 + \beta_1 \times Detection_{ijk})}$<br><br>must be applied to estimate the proportion of plasticine models attacked or not attacked using the model's estimated parameters. | models have a different status in the e.g., dataset (2) in comparison to dataset (1). This means that 8% of attacked models are of the same status, regardless of the detection type.<br><br>We used pakage *simr* (Green et al. 2023) to perform power analyses on each of these 3 datasets.<br><br>Power analyses indicate a result of over 99% for the first two data sets (see Fig.S2). This suggests a very high probability of detecting significant differences in our sample size, if any. On the other hand, if the predation clues assessed by human observers accurately reflect real predation (predation evidence), the power analysis drops drastically to 0%. This remains logical, since in this case there is absolutely no difference according to the type of detection. |

| | | | | | | |
|---|---|---|---|---|---|---|
| 3. Does the predation on corpse vs. plasticine model differ? | H3. Given birds' ability to recognize fake and real prey, the number of attacks on corpses is higher compared to the models, both assessed by camera traps. | (H3) Camera traps will be used to assign the status "attacked" or "not attacked" by a bird to each of the 420 corpses and 420 models. | $logit(P(Y_{ij} = 1)) = \beta_0 + \beta_1 \times PreyType_{Models,ij} + \gamma_j$<br><br>where $Y_{ij}$ is the binary variable, indicating the presence or absence of bird predation at each branch (one data point per prey type and branch). Specifically, $Y_{ij}$=1 indicates at least one instance of bird predation evidence, and $Y_{ij}$=0 indicates no evidence of bird predation. $\beta_0$ is the model intercept, representing the logit-transformed probability of predation on the corpses (i.e., $PreyType_{Corpse,ij}$), $\beta_1$ the coefficient of the fixed effect of plasticine models (i.e., $PreyType_{Model,ij}$). $\gamma_j$ is the random effect for tree identity nested in temporal permutation and follows a normal law $\gamma_j$~N(0,$\sigma_\gamma^2$).<br><br>We predict $\beta_1 < 0$ because the average level of caterpillars attacked would be lower with the "model" treatment than with the "corpse" treatment.<br><br>The mathematical transformation | In the same way as for the previous hypothesis, we have created 3 different datasets with different effect sizes:<br><br>(1) Attack rate of 5% for models versus 10% for corpses; (2) 5% models vs. 20% corpses; (3) 10% models vs. 20% corpses.<br><br>We performed power analyses on each of these 3 datasets.<br><br>The 3 power analyses with 1000 simulations indicate a statistical power of over 75% for the 3 different scenarios (see Fig. S3). This suggests that our sample size has a high probability of detecting significant differences if they really exist. | If $\beta_1 \simeq 0$, there is approximately the same level of attack on corpses and on models, i.e., birds do not distinguish between the two types of caterpillars.<br><br>If $\beta_1 > 0$, the average level of caterpillars attacked would be higher on models than on corpses, i.e., the predation attempts done on models do not reflect the predation. | If predation attempts on models do not reflect predation as observed on corpses, this would argue for the use of corpses in studies of predation interactions between insectivorous birds and herbivorous caterpillars. However, we would need to be cautious about the species used, and carefully discuss their condition (live or corpse) at the time of experimentation. |

| | | | | | | |
|---|---|---|---|---|---|---|
| | | | $p = \frac{\exp(\beta_0+\beta_1 \times PreyType_{ij})}{1+\exp(\beta_0+\beta_1 \times PreyType_{ij})}$ must be applied to estimate the proportion of caterpillars attacked or not attacked using the model's estimated parameters. | | | |
| 4. Is there a bias between observer and camera detection of bird marks on corpses?<br><br>*If no, and according to the results for H3, we might be able to answer :*<br><br>Are real caterpillars a more accurate method than plasticine caterpillars for quantifying predation of herbivorous insects by birds if not | H4. The observer detection of predation clues leads to an overestimation of the predation on corpses accounting for missing individuals as a predation event. | (H4) Bird predation evidence will be recorded on each of the 420 corpses using camera traps. In the field, we will assess bird predation clues on each corpses. A missing corpse or with a missing part will be considered attacked. Each corpse will therefore be considered attacked or not attacked by 2 types of detection (clues or evidence). | Exactly the same model as for hypothesis H2 on real dead caterpillars.<br>$logit(P(Y_{ijk} = 1)) = \beta_0 + \beta_1 \times Detection_{Clues,ijk} + \gamma_j + \zeta_k$<br>where $Y_{ijk}$ is the binary response variable (one data point per detection method, branch, and model), with $Y_{ijk}$=1 indicating at least one predation clue or one bird presence evidence, and $Y_{ijk}$=0 indicating no detection. $\beta_0$ is the model intercept, representing the logit-transformed probability of detection for the camera-assessed bird presence evidence (i.e., $Detection_{Evidence,ijk}$), $\beta_1$ the coefficient of the fixed effect predation clues detected by the observer (i.e., $Detection_{Clues,ijk}$). $\gamma_j$ is the | The power analyses of hypothesis H2 are completely transposable to this hypothesis (see Fig. S2). | If $\beta_1 \simeq 0$, predation clues exactly reflect predation evidence assessed by the camera.<br><br>If $\beta_1 < 0$, predation clues are underestimated in relation to predation evidence assessed by the cameras. | In the case where $\beta_1 \simeq 0$, and only in this case, there is no need for camera trap detection to assess predation on corpses; detection by human observers is sufficient. We can then refer to hypothesis H3 to determine whether corpses are indeed a better method for assessing bird predation on herbivorous insects than plasticine models. |

| | | | |
|---|---|---|---|
| complemented with the detection of predation evidence assessed by camera traps? | | | random effect for tree identity nested in temporal permutation and follows a normal law $\gamma_j \sim N(0,\sigma_\gamma^2)$, $\zeta_k$ is the random effect for corpse identity and follows a normal law $\zeta_k \sim N(0,\sigma_\zeta^2)$.<br><br>We predict $\beta_1 > 0$ because it would reflect additional predation clues compared to predation evidence. |

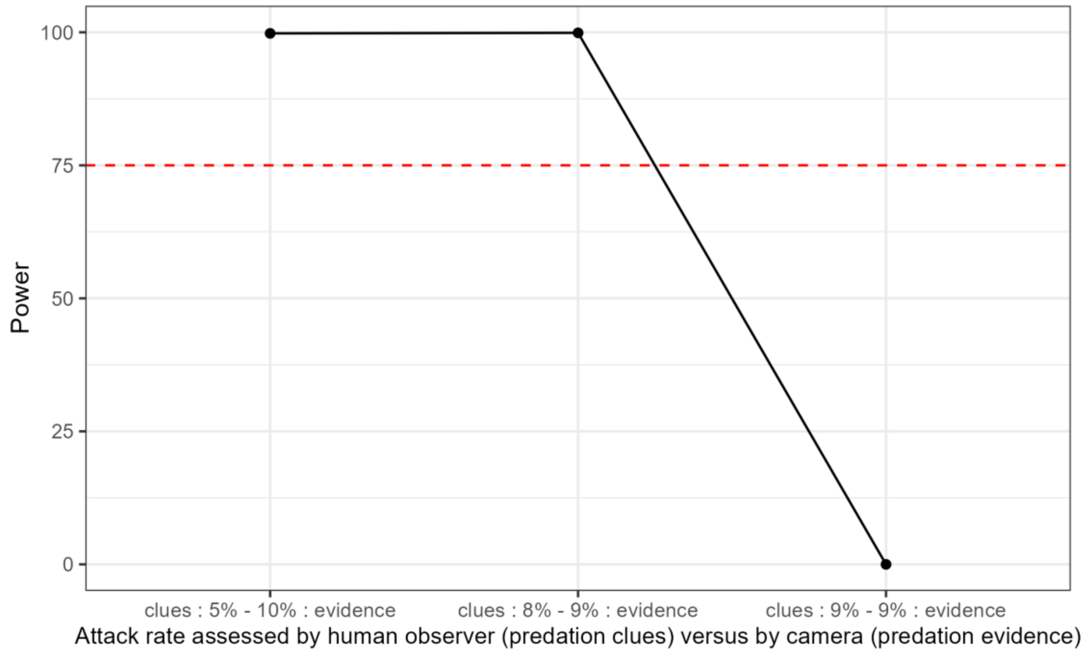

**S2 Fig. The results of power analyses with 1000 simulations for the three different scenarios tested (hypotheses 2 and 4).**

The power analysis results according to the three different datasets are the following: predation clues 5% - 10% predation evidence: 99.80% (range: 99.28, 99.98); clues 8% - 9% evidence: 99.90% (range: 99.44, 100.00); clues 9% - 9% evidence: 0.00% (range: 0.00, 0.37).

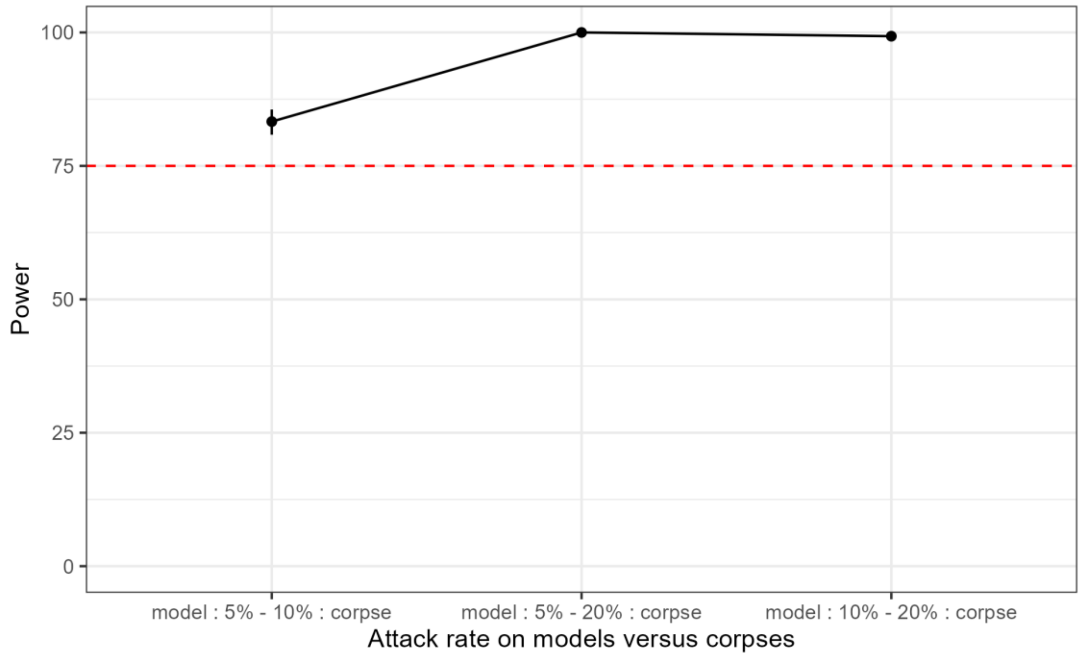

**S3 Fig. The results of power analyses with 1000 simulations for the 3 different scenarios tested (hypothesis 3).**

The power analyses were based on the sample size of 420 caterpillars per fixed factor. The power analysis results according to the three different datasets are the following: model 5% - 10% corpse: 83.30% (range: 80.84, 85.56); model 5% - 20% corpse: 100.00% (range: 99.63, 100.00); model 10% - 20% corpse: 99.30% (range: 98.56, 99.72).

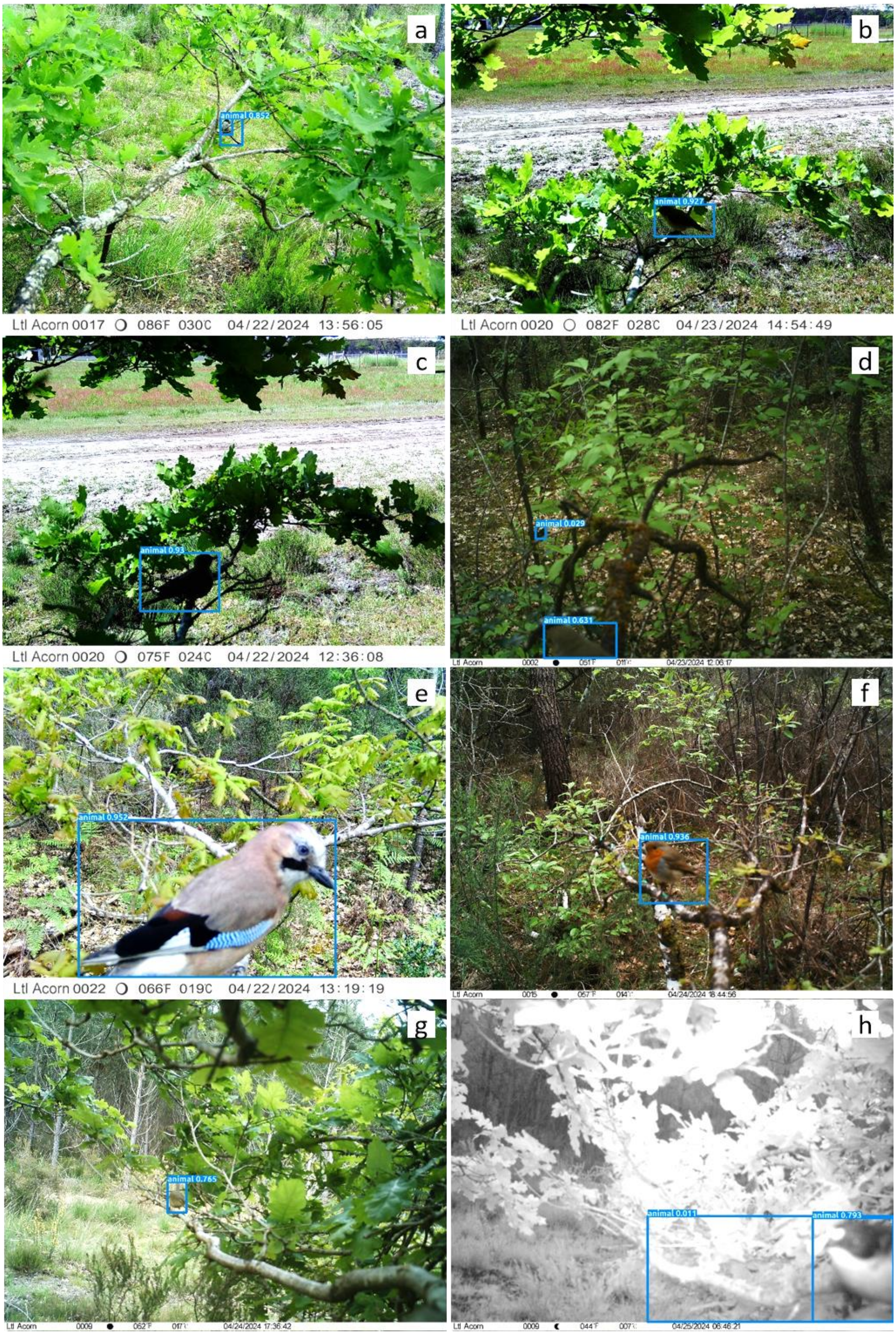
a
animal 0.852
Ltl Acorn 0017 086F 030C 04/22/2024 13:56:05
b
animal 0.927
Ltl Acorn 0020 082F 028C 04/23/2024 14:54:49
c
animal 0.93
Ltl Acorn 0020 075F 024C 04/22/2024 12:36:08
d
animal 0.029
animal 0.631
Ltl Acorn
04/23/2024 12:06:17
e
animal 0.952
Ltl Acorn 0022 066F 019C 04/22/2024 13:19:19
f
animal 0.936
Ltl Acorn
04/24/2024 18:44:56
g
animal 0.765
Ltl Acorn
04/24/2024 17:36:42
h
animal 0.011
animal 0.793
Ltl Acorn
04/25/2024 06:46:21

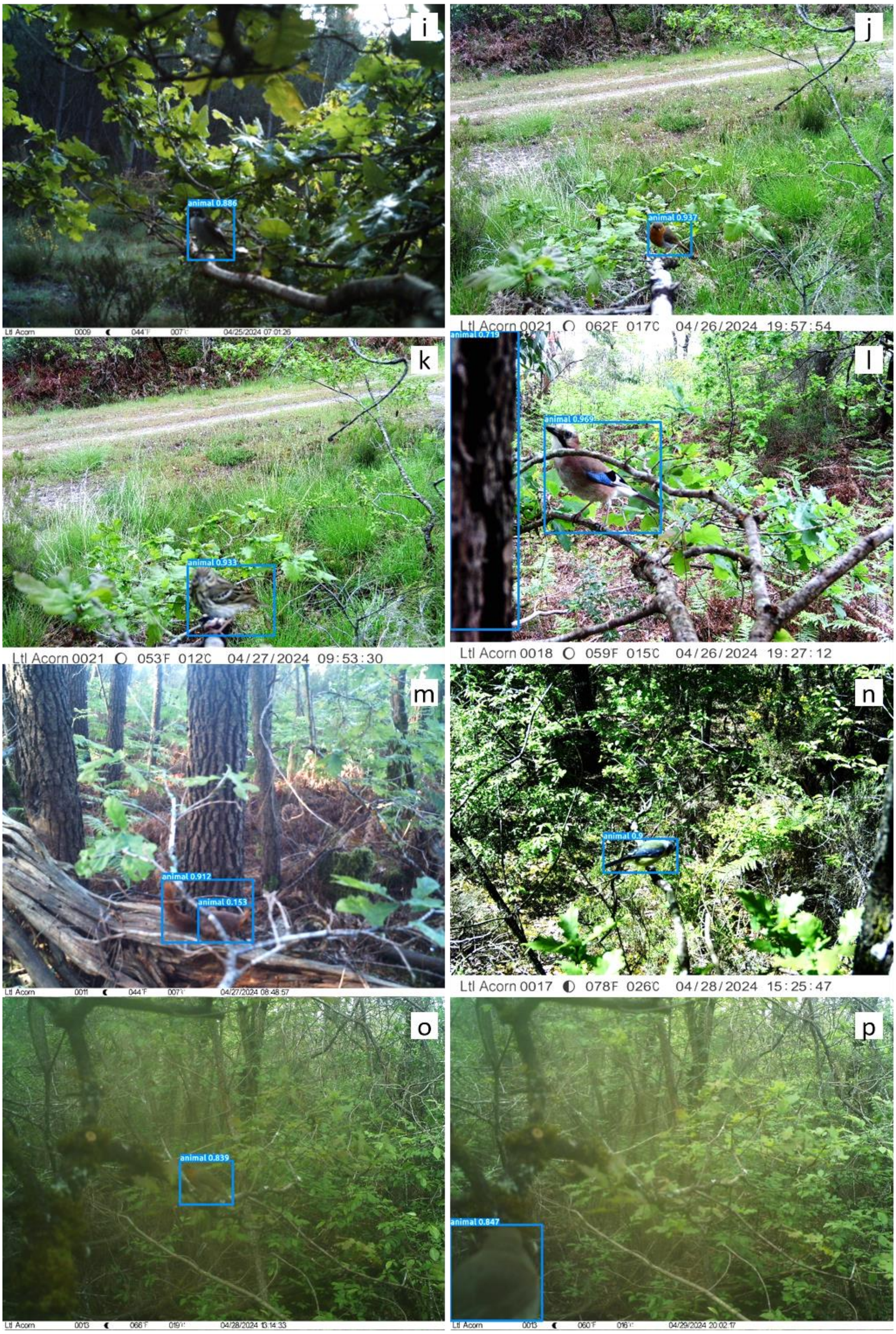

i
animal 0.886
j
animal 0.937
Ltl Acorn 0021 062F 017C 04/26/2024 19:57:54
k
animal 0.933
Ltl Acorn 0021 053F 012C 04/27/2024 09:53:30
l
animal 0.719
animal 0.969
Ltl Acorn 0018 059F 015C 04/26/2024 19:27:12
m
animal 0.912
animal 0.153
n
Ltl Acorn 0017 078F 026C 04/28/2024 15:25:47
o
animal 0.839
p
animal 0.847

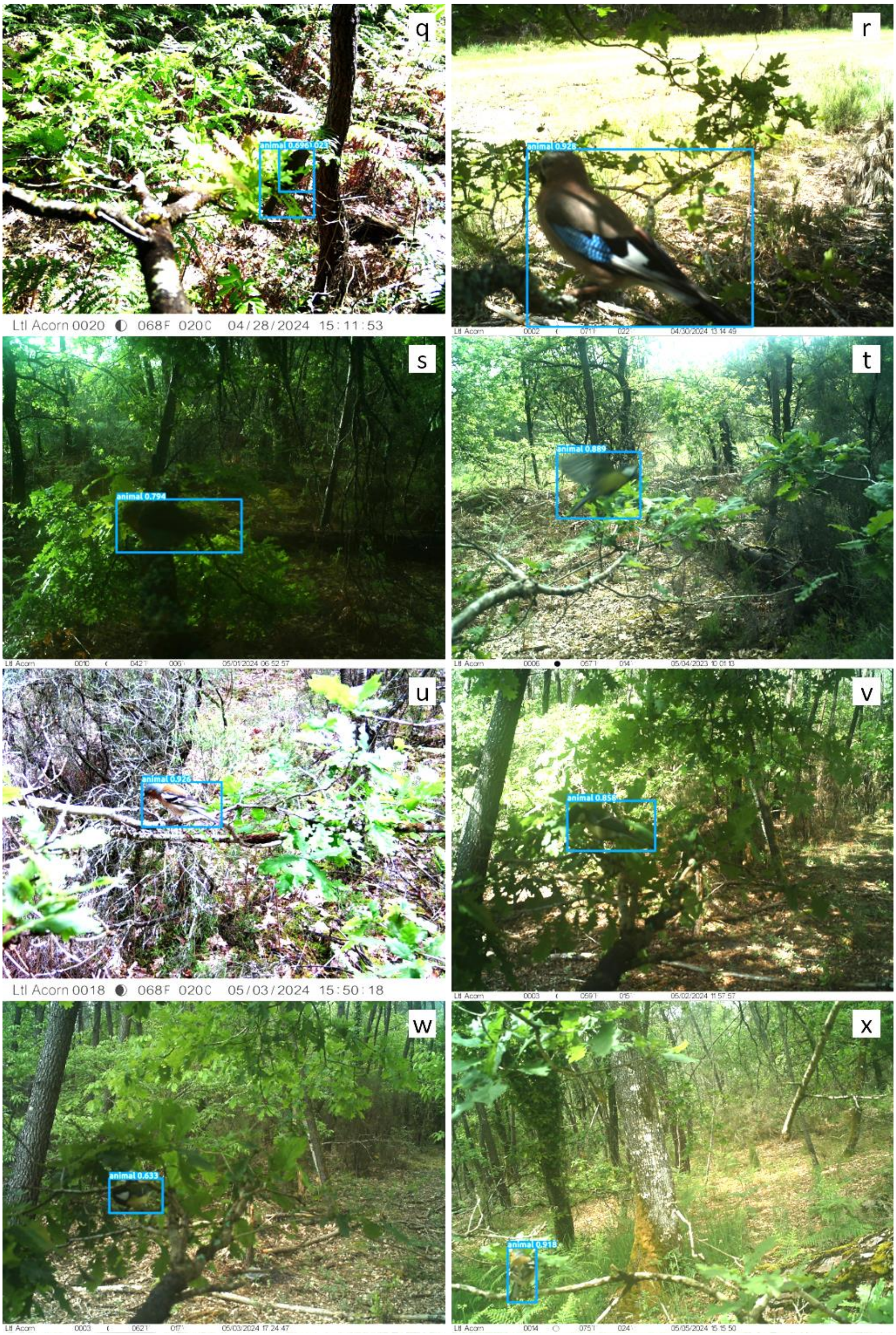
q
animal 0.696
Ltl Acorn 0020 068F 020C 04/28/2024 15:11:53
r
animal 0.928
s
animal 0.794
t
animal 0.889
u
animal 0.926
Ltl Acorn 0018 068F 020C 05/03/2024 15:50:18
v
animal 0.858
w
animal 0.633
x
animal 0.918

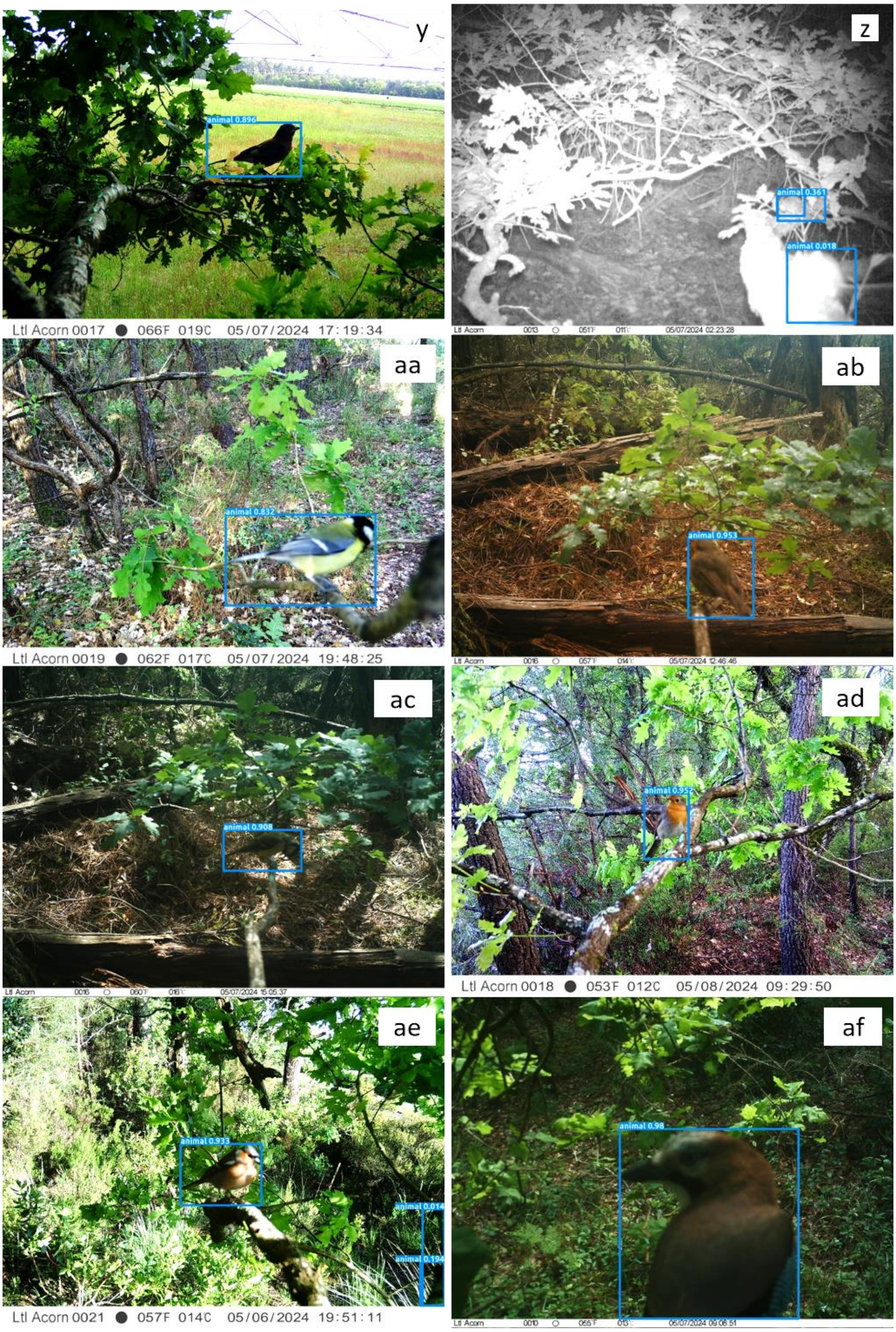
y
animal 0.896
Ltl Acorn 0017 066F 019C 05/07/2024 17:19:34
z
animal 0.361
animal 0.018
aa
animal 0.832
Ltl Acorn 0019 062F 017C 05/07/2024 19:48:25
ab
animal 0.953
ac
animal 0.908
ad
animal 0.952
Ltl Acorn 0018 053F 012C 05/08/2024 09:29:50
ae
animal 0.933
animal 0.014
animal 0.194
Ltl Acorn 0021 057F 014C 05/06/2024 19:51:11
af
animal 0.98

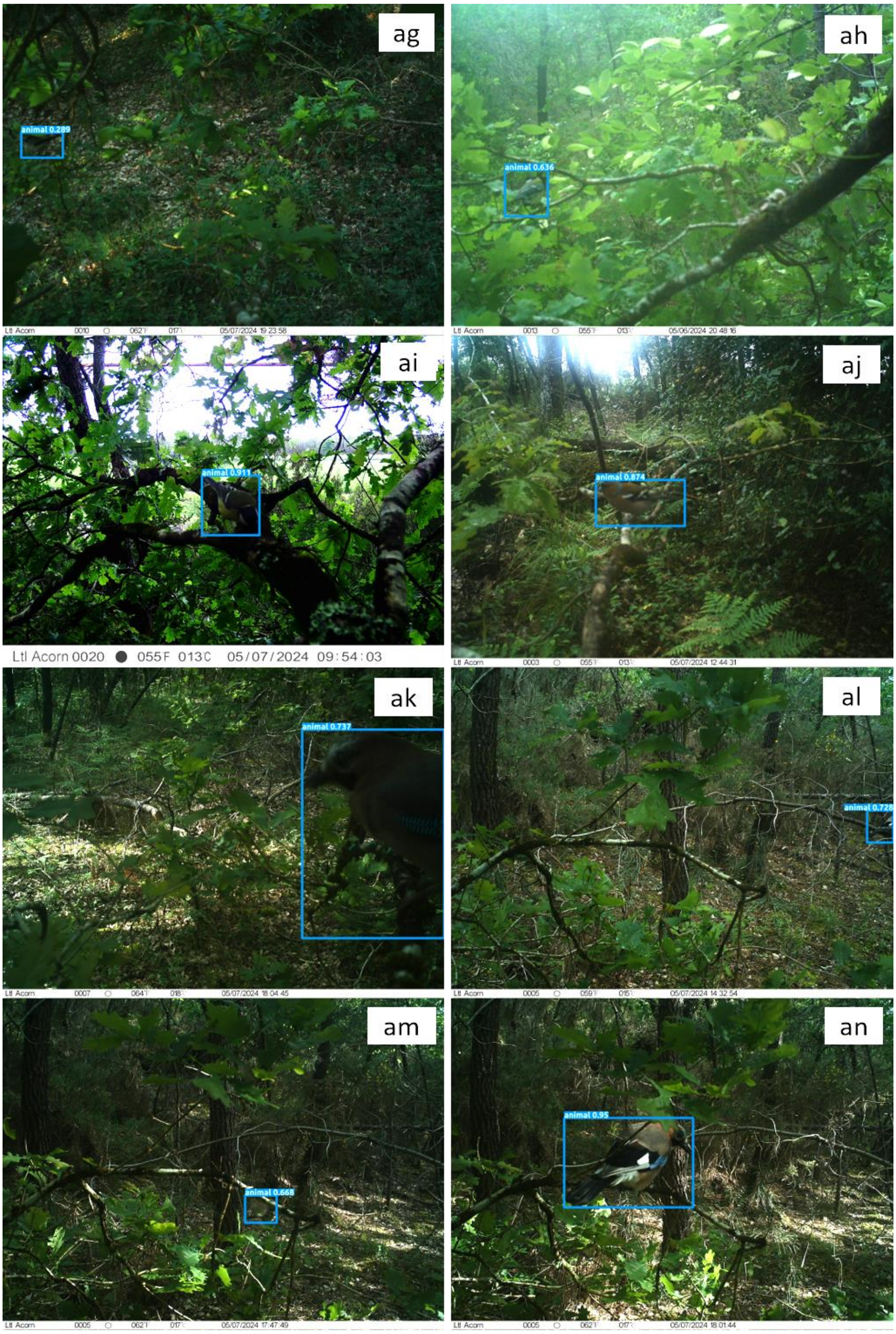
ag
animal 0.289
ah
animal 0.636
ai
animal 0.911
Ltl Acorn 0020 055F 013C 05/07/2024 09:54:03
aj
animal 0.874
ak
animal 0.737
al
animal 0.728
am
animal 0.668
an
animal 0.95

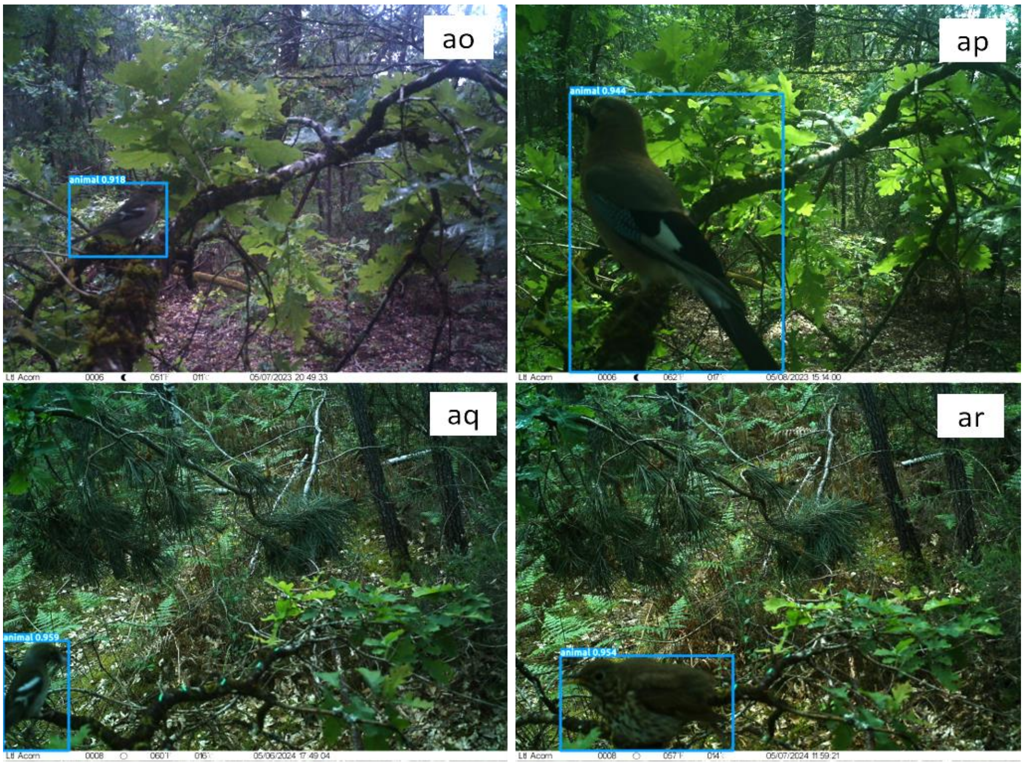

**S4 Fig. All detected birds by MegaDetector AI software.**
Camera traps photographs of potential predators of caterpillars. The blue squares correspond to MegaDetector detections with the associated confidence level. The animal species included are: the Long-tailed Tit (*Aegithalos caudatus*, S1a and S1ag Figs), the Tree Pipit (*Anthus trivialis*, S1c and S1k Figs), the Common Chiffchaff (*Phylloscopus collybita*, S1b and S1g Figs), the European Robin (*Erithacus rubecula*, S1d, S1f, S1j, S1o, S1p, S1q, S1s, S1ab, and S1ad Figs), the Eurasian Jay (*Garrulus glandarius*, S1e, S1l, S1r, S1y, S1af, S1ak, S1an, and S1ap Figs), the European Pied Flycatcher (*Ficedula hypoleuca*, S1h and S1i Fig.), the Red Squirrel (*Sciurus vulgaris*, S1m Fig), the Eurasian Blue Tit (*Cyanistes caerulus*, S1n and S1am Figs), the Great Tit (*Parus major*, S1t, S1v, S1w, S1aa, S1ac, S1ah and S1ai Figs), the Common Chaffinch (*Fringilla coelebs*, S1u, S1x, S1ae, S1aj, S1al, S1ao and S1aq Figs), unidentified rodents (S1z Fig), and the Song Trush (*Turdus philomelos*, S1ar Fig).

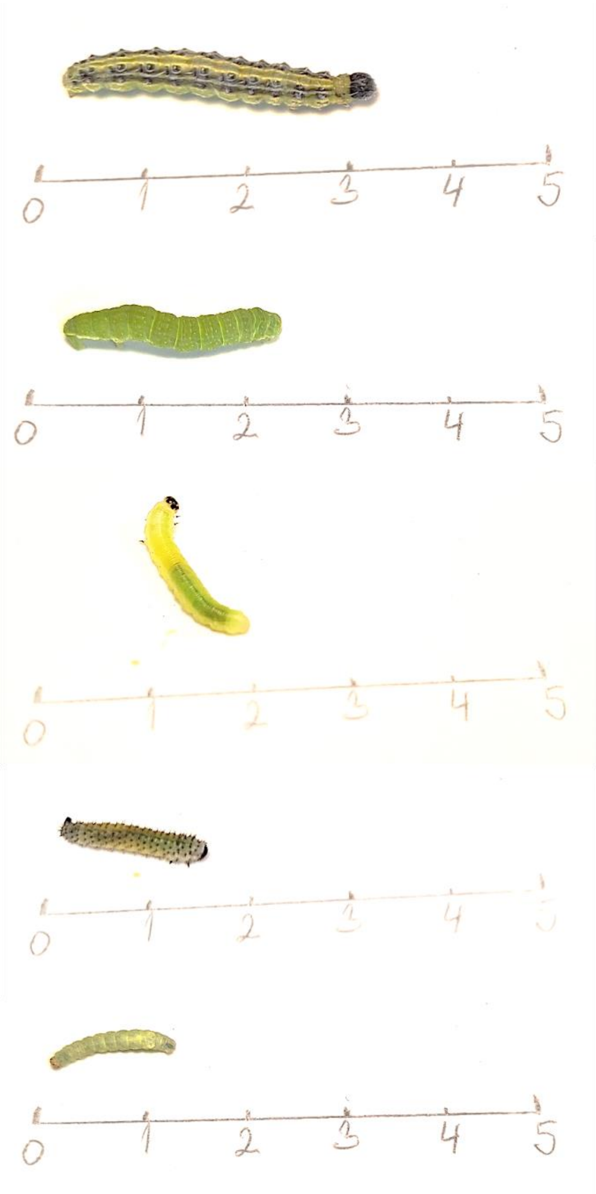

**S5 Fig. Photographs of larvae found in the study area.**

From top to bottom, the families, genera, or species are: *Cydalima perspectalis* (non-native species), Geometridae family, Tenthredinidae family (likely of Caliroa genus), Tenthredinidae family (likely of Periclista genus), *Operopthera brumata*. Scale in cm.
